# Robust regulatory architecture of pan-neuronal gene expression

**DOI:** 10.1101/2021.12.10.472091

**Authors:** Eduardo Leyva-Díaz, Oliver Hobert

## Abstract

Pan-neuronally expressed genes, such as genes involved in the synaptic vesicle release cycle or in neuropeptide maturation, are critical for proper function of all neurons, but the transcriptional control mechanisms that direct such genes to all neurons of a nervous system remain poorly understood. We show here that six members of the CUT family of homeobox genes control pan-neuronal identity specification in *C. elegans*. Single CUT mutants show barely any effects on pan-neuronal gene expression or global nervous system function, but such effects become apparent and progressively worsen upon removal of additional CUT family members, indicating a critical role of gene dosage. Overexpression of each individual CUT gene rescued the phenotype of compound mutants, corroborating that gene dosage, rather than the activity of specific members of the gene family, is critical for CUT gene function. Genome-wide binding profiles as well as mutation of CUT binding sites by CRISPR/Cas9 genome engineering show that CUT genes directly control the expression of pan-neuronal features. Moreover, CUT genes act in conjunction with neuron-type specific transcription factors to control pan-neuronal gene expression. Our study, therefore, provides a previously missing key insight into how neuronal gene expression programs are specified and reveals a highly buffered and robust mechanism that controls the most critical functional features of all neuronal cell types.

## INTRODUCTION

To understand nervous system development, it is of critical importance to decipher the mechanisms that control the expression of neuronal gene batteries. Apart from ubiquitous housekeeping genes expressed in all tissue types, neuronal gene batteries fall into two categories: (1) Genes selectively expressed in specific neuron classes; these include neurotransmitter synthesis pathway genes, individual neuropeptides genes, ion channels, signaling receptors and many others (**Figure 1A**)(Hobert et al., 2010; Taylor et al., 2021). (2) Pan-neuronally expressed genes that execute functions shared by all neurons, but not necessarily other cell types; these genes encode proteins involved in a number of generic neuronal processes, including synaptic vesicle release (e.g., *RAB3, SNAP25, RIM*), dense core biogenesis and release (e.g., *CAPS*), molecular motors (e.g. kinesins) or neuropeptide processing enzymes (e.g., endo- and carboxypeptidases, monooxygenases)(Stefanakis et al., 2015; Taylor et al., 2021). Great strides have been made in understanding the regulation of the first category of genes, neuron type-specific gene batteries, in the nervous system of many species (Hobert, 2016; Hobert and Kratsios, 2019; Qiu and Ghosh, 2008). However, the regulatory programs that orchestrate pan-neuronal gene expression have remained elusive in any species to date (Stefanakis et al., 2015). Proneural transcription factors of the basic helix-loop-helix family control pan-neuronal gene batteries only indirectly since their expression is transient and fades either before terminal differentiation or during the transcriptional maintenance of neuronal gene batteries in juvenile and adult life (Bertrand et al., 2002).

**Figure 1.**
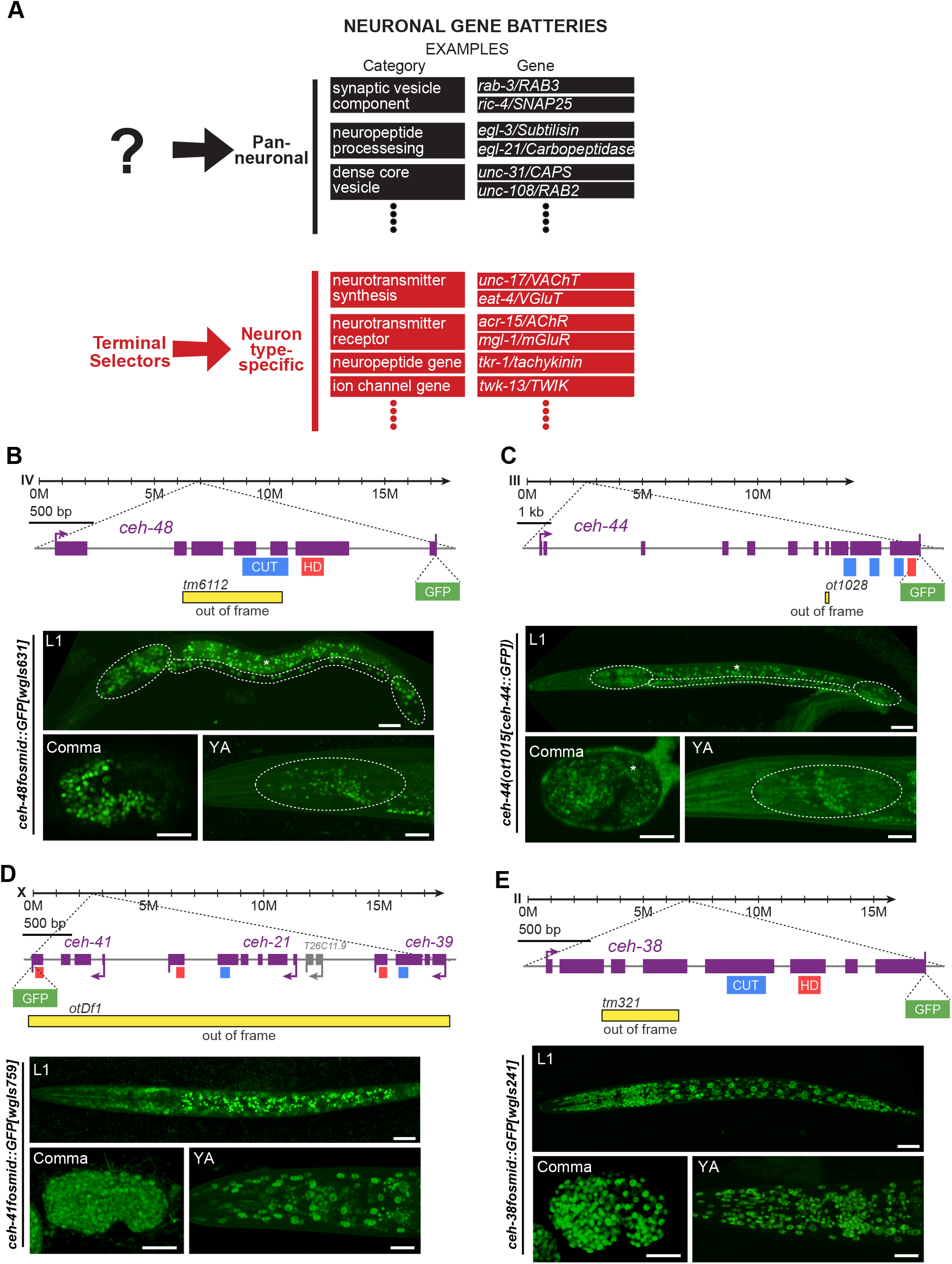
CUT genes are expressed pan-neuronally. **(A)** Schematic illustration of two main components of neuronal gene batteries, pan- neuronally expressed genes, for which no current regulator is known, and neuron type- specific gene batteries that are controlled by terminal selector-type transcription factors (Hobert et al., 2010). **(B-E)** Schematic representation of *ceh-48* (B), *ceh-44* (C), *ceh-41*, *ceh-21* and *ceh-39* (D), and *ceh-38* (E), gene loci showing mutant alleles, GFP tags, and CUT and Homeodomain motifs location. Reporter expression at the comma embryonic stage (bottom left, lateral view), L1 larval stage (top, full worm lateral views) and young adult stage (bottom right, lateral view of the head) showing *ceh-48* (*ceh- 48fosmid::GFP[wgIs631]*) (B) and *ceh-44* (*ceh-44(ot1015[ceh-44::GFP])*) (C) pan- neuronal expression, and *ceh-41*, *ceh-21* and *ceh-39* (D), *ceh-38* (*ceh- 38fosmid::GFP[wgIs241]*) (E) ubiquitous expression. We use a fosmid reporter for *ceh- 41* (*ceh-41fosmid::GFP[wgIs759]*), the last gene in the operon of three ONECUT genes, which provides a read-out for expression of all genes in the operon. The embryonic comma stage is the stage when neurons are born. Head ganglia, ventral nerve cord, and tail ganglia outlined in L1 images, and head ganglia outlined in YA images for *ceh-48* and *ceh-44* reporters. Asterisks (*) indicate autofluorescence in L1 (*ceh-48* and *ceh-44*) and Comma (*ceh-44*) images. Comparison of the expression level of all *gfp* CUT gene loci, assessed in newly generated CRISPR/Cas9-engineered reporter alleles, shows that *ceh- 38* is the highest expressed CUT family member (**Fig. S1A**). Scale bars 15 μm.

In the nematode *Caenorhabditis elegans,* and other organisms as well, the expression of neuron type-specific genes during terminal differentiation is controlled by neuron-type specific combinations of terminal selector transcription factors (Hobert, 2016; Hobert et al., 2010; Hobert and Kratsios, 2019). However, genetic removal of a terminal selector does not generally affect the expression of pan-neuronal identity features (Hobert, 2016; Hobert et al., 2010). For example, loss of the LIM homeobox gene *ttx-3* or the EBF-type *unc-3* zinc knuckle transcription factor results in the loss of all known neuron type-specific identity features of the cholinergic AIY interneuron or the cholinergic ventral nerve cord motorneurons, respectively, while the expression of pan-neuronal genes remains unaffected (Altun-Gultekin et al., 2001; Kratsios et al., 2011). Similarly, in mice, BRNA3 and ISL1 control neuron type-specific, but not pan-neuronal features of sensory neurons of the trigeminal ganglion and dorsal root ganglia (Dykes et al., 2011). In attempts to decipher the apparent parallel acting gene regulatory programs of pan-neuronal gene expression, we have previously isolated a multitude of *cis-*regulatory enhancer elements from pan-neuronally expressed genes (Stefanakis et al., 2015). However, genetic screens for mutants that affect expression of these *cis*-regulatory elements have remained unsuccessful (Leyva-Diaz et al., 2017).

In this paper, we describe the discovery that all members of a subfamily of homeobox genes, the CUT homeobox genes, jointly control pan-neuronal gene expression. CUT genes are expressed in all neurons and bind to the regulatory control regions of pan-neuronal genes. Deletion of the CUT binding motif from pan-neuronal genes, using CRISPR/Cas9 genome engineering, disrupts expression and function of pan-neuronal genes. Removal of individual CUT genes reveals a dosage-sensitive function of these genes in controlling pan-neuronal gene expression and neuronal function. These phenotypes can be rescued by the expression of individual CUT factors, indicating that these factors act redundantly. A more extensive neuronal transcriptional profiling in neurons lacking all neuronal CUT genes reveals that these factors are required for the expression of large cohorts of neuronal genes. Further genetic loss of function analysis reveals that pan-neuronally expressed CUT genes cooperate with neuron type-specific terminal selectors to control pan-neuronal gene expression. Our studies reveal an exceptionally robust regulatory architecture of pan-neuronal gene expression, contrasting the regulation of neuron-type specific gene regulations. Our findings may have implications for the evolution of neuronal cell type diversity.

## RESULTS

### Cut homeobox genes are expressed in all neurons

Our recently reported genome-wide analysis of the expression of all homeobox genes, critical regulators of neuron-type specific identity programs (Reilly et al., 2020), uncovered a clue for potentially solving the riddle of pan-neuronal gene expression. Using both fosmid-based reporters as well as CRISPR/Cas9-engineered reporter alleles, in which we inserted *gfp* reporter transgenes in endogenous gene loci, we found that two homeobox genes, *ceh-44* and *ceh-48,* are restricted to all neurons of the adult nervous system (**Figure 1B and 1C**). The only non-neuronal cells that express one of these two genes (*ceh-48)* are the secretory uv1 uterine cells, whose neuronal characters, including expression of synaptic vesicular machinery and the neurotransmitter tyramine, have been noted before (Alkema et al., 2005; Stefanakis et al., 2015). Expression of *ceh-44* and *ceh- 48* commences right after the birth of neurons in the embryo, slightly preceding the onset of various other markers of pan-neuronal identity (Stefanakis et al., 2015), and they are continuously expressed throughout the life of the organism (**Figure 1B and 1C**).

*ceh-44* and *ceh-48* are members of the CUT family of homeobox genes, defined by the presence of a homeodomain and one or more CUT domains (Burglin and Affolter, 2016). Based on the presence of multiple CUT domains, *ceh-44* is the sole representative of the CUX subclass of the CUT family in *C. elegans*, while *ceh-48* is a member of the ONECUT subclass, characterized by the presence of a single CUT domain (Burglin and Cassata, 2002). The *C. elegans* genome encodes five additional ONECUT genes, three of which are located in a single operon (**Figure 1D**). While *ceh-48* is pan-neuronally expressed, four of these additional ONECUT genes are ubiquitously expressed in all tissues at all stages (**Figure 1D and 1E**), while one ONECUT gene (*ceh-49*) is only expressed in the early embryo before neurogenesis. *ceh-49* was not considered further here. Comparison of the expression level of all CUT gene loci, assessed with CRISPR/Cas9-engineered reporter alleles, shows that *ceh-38* is the highest expressed CUT family member (**Figure S1A**).

### Binding sites for Cut homeodomain proteins are required for pan-neuronal gene expression

The pan-neuronal expression of *ceh-44* and *ceh-48* made us consider these CUT family genes as potential regulators of pan-neuronal identity. Supporting this notion we find that: (a) the many pan-neuronal genes whose *cis*-regulatory control regions we had previously defined to contribute to pan-neuronal gene expression contain predicted CUT homeodomain binding sites; and that (b) animal-wide chromatin immunoprecipitation (ChIP) of CEH-48 from the modENCODE project reveals binding of CEH-48 to these *cis-* regulatory elements (**Figures 2A** and S2A; **Table S1**) (Davis et al., 2018; Stefanakis et al., 2015). We assessed the functional relevance of these CUT binding sites in two different ways: First, we deleted these sites in the context of enhancer fragments, isolated from pan-neuronal gene loci, that drive broad neuronal if not pan-neuronal expression in transgenic, multicopy reporter arrays. We found a severe loss of expression upon deletion of CUT sites from isolated *cis*-regulatory enhancer elements derived from the *rab- 3/RAB3*, *ric-4/SNAP25* and *unc-10/RIM* genes (**Figure 2B-E**). Second, we used CRISPR/Cas9 genome engineering to first tag several pan-neuronal genes (*rab-3/RAB3, ric-4/SNAP25, unc-10/RIM, ehs-1/EPS15*) with a *gfp* reporter tag, and to subsequently delete their respective CUT homeodomain binding site from the respective endogenous locus. Deletion of CUT homeodomain binding sites affected expression of all four pan-neuronal genes that we tested (**Figure 2B-E and S5D**).

**Figure 2.**
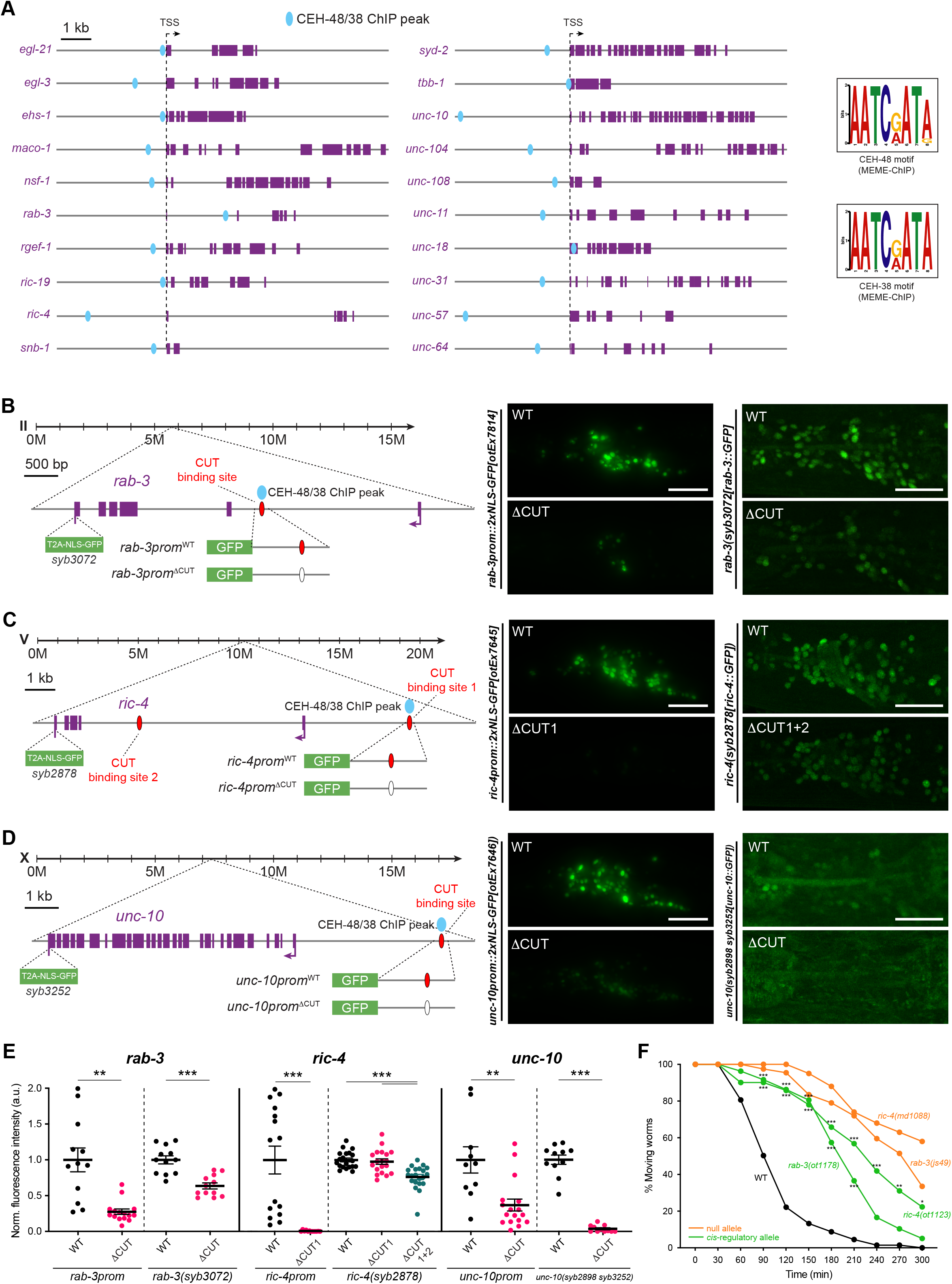
CUT genes binding is required for pan-neuronal gene expression. **(A)** Schematic representation of 20 pan-neuronal genes and the location of CEH-48 and CEH-38 peaks found in the ChIP-seq datasets. CEH-48 and CEH-38 peaks overlap for all genes except in *maco-1* and *tbb-1*, which only contain CEH-38 peaks, and *ric-19*, which only contains a CEH-48 peak. Scale represents 2kb for *unc-104* and *unc-31*. The consensus binding motifs for CEH-48 and CEH-38, extracted from the ChIP-seq datasets using MEME-ChIP (Machanick and Bailey, 2011) are shown on the right. See **Table S1** for a full list of genes with CEH-48 and CEH-38 ChIP peaks. See **Figure S2** for how CUT ChIP binding correlates with the *cis-*regulatory elements that we previously defined in pan-neuronally expressed genes (Stefanakis et al., 2015). **(B-D)** Schematic representation of *rab-3* (B), *ric-4* (C) and *unc-10* (D) gene loci (left) showing the location of CEH-48/CEH-38 ChIP peaks, CUT binding sites, endogenous GFP tags for CRISPR reporters (*rab-3(syb3072[rab-3::T2A::3xNLS::GFP])*, *ric- 4(syb2878[ric-4::T2A::3xNLS::GFP])*, *unc-10(syb2898 syb3252[unc- 10::T2A::3xNLS::GFP])*), and small promoters tested (*rab-3prom10::2xNLS- GFP[otEx7814]*, *ric-4prom30::2xNLS-GFP[otEx7645]*, *unc-10prom12::2xNLS- GFP[otEx7646]*). Blue ovals indicate binding based on ChIP-seq peak data, red ovals indicate binding site based on sequence. Worm head GFP images showing a reduction in pan-neuronal gene expression when the CUT binding site is mutated compared to WT (middle, left). Mutation of the same CUT binding sites endogenously in the context of CRISPR reporters affects pan-neuronal expression (middle, right). *ric-4 gfp*-tagged allele expression is only affected upon mutation of additional CUT binding sites (site 1 and 2). *unc-10 gfp*-tagged allele expression is very dim and expression is not visible in all neurons. All images correspond to worms at the L4 larval stage. **(E)** Quantification of small promoters and CRISPR reporters (shown in B-D) head neurons fluorescence intensity in wild-type and upon CUT site mutations in the regulatory control regions of *rab-3* (left), *ric-4* (center) and *unc-10* (right). The data are presented as individual values with each dot representing the expression level of one worm with the mean ± SEM indicated. Unpaired *t*-test, **P < 0.01, ***P < 0.001. For *ric-4(syb2878[ric-4::T2A::3xNLS::GFP])*, one-way ANOVA followed by Tukey’s multiple comparisons test; ***P < 0.001. n ≥ 10 for all genotypes. **(F)** Aldicarb-sensitivity defects in wild-type animals, *ric-4* and *rab-3 cis*-regulatory alleles (*ric-4(ot1123 syb2878)*, *rab-3(ot1178 syb3072)*), and *ric-4* and *rab-3* null alleles (*ric- 4(md1088)*, *rab-3(js49)*). Wild-type data is represented with black dots, the *cis*-regulatory alleles with green dots, and null alleles with orange dots. Two-way ANOVA followed by Tukey’s multiple comparisons test, comparisons for *ric-4* and *rab-3 cis*-regulatory alleles vs wild-type indicated; *P < 0.05, **P < 0.01, ***P < 0.001. n = 3 independent experiments (25 animals per independent experiment). TSS, transcription start site; WT, wild-type; a.u., arbitrary units. Scale bars 15 μm for all panels except for CRISPR reporters in (B-D), where scale bars equal 10 μm.

We tested the functional significance of the CUT site mutations by asking whether these potential *cis*-regulatory alleles displayed behavioral defects expected from the loss of function of these pan-neuronal genes. *rab-3/RAB3* and *ric-4/SNAP25* null alleles show defects in synaptic transmission that can be measured via the sensitivity of animals to a drug that affects synaptic transmission at the neuromuscular junction, aldicarb (Nguyen et al., 1995; Nonet et al., 1997). We found that *rab-3/RAB3* and *ric-4/SNAP25* alleles carrying CUT site mutations show resistance to aldicarb (**Figure 2F**), which correlates with the reduction in *ric-4/SNAP25* and *rab-3/RAB3* expression observed in these alleles, and indicate impairment on synaptic transmission. Taken together, the functional relevance of presumptive CUT homeodomain binding sites hints toward a function of the CUT family of transcription factors as potential regulators of pan-neuronal gene expression.

### Dosage-dependent requirement of Cut homeobox genes for pan-neuronal gene expression and neuronal behavior

We next analyzed the consequences of genetic removal of the two pan-neuronally expressed *ceh-44* and *ceh-48* genes. We used a *ceh-48* null allele from a *C. elegans* knockout consortium and engineered a *ceh-44* null allele using the CRISPR/Cas9 system. As a first step to assess gene function, we analyzed the expression of a *rab-3* reporter construct in single and double *ceh-44* and *ceh-48* null mutant backgrounds. Given the functional importance of the CUT site in the *rab-3* locus described above, we were surprised to observe no *rab-3/RAB3* expression defects in either single or *ceh-44; ceh- 48* double null mutant animals (**Figure 3A**).

**Figure 3.**
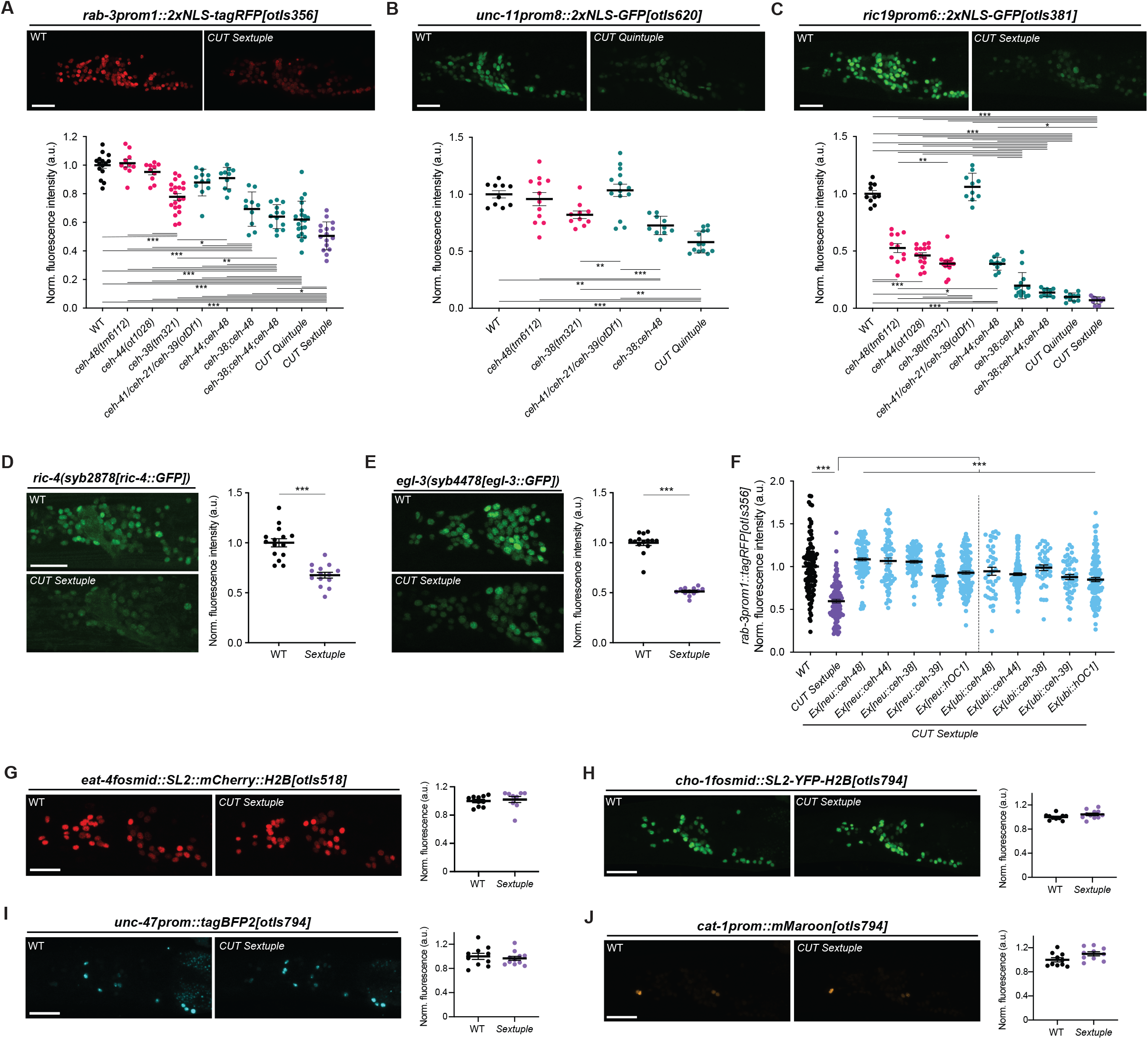
CUT genes act in a dosage-dependent manner to control pan-neuronal gene expression. **(A-C)** Expression of *rab-3prom1::2xNLS-tagRFP[otIs356]* (A), *unc- 11prom8::2xGFP[otIs620]* (B) and *ric19prom6::2xNLS-GFP[otIs381]* (C) in wild-type (left) and CUT sextuple mutant (right). Lateral views of the worm head at the L4 stage are shown. Quantification of fluorescence intensity in head neurons (bottom) in wild-type, individual CUT mutants (*ceh-48(tm6112)*, *ceh-44(ot1028)* and *ceh-38(tm321)*) and compound CUT mutants (*otDf1*, which deletes *ceh-41*, *ceh-21* and *ceh-39*; double *ceh- 44;ceh-48*, double *ceh-38;ceh-48*, triple *ceh-38;ceh-44;ceh-48*, quintuple *ceh-38;ceh- 48;otDf1*, and sextuple *ceh-38;ceh-44;ceh-48;otDf1*). *unc-11prom::2xGFP[otIs620]* and *ceh-44* are located in the same chromosome (chr. I) and cannot be recombined together. The data are presented as individual values with each dot representing the expression level of one worm with the mean ± SEM indicated. Wild-type data is represented with black dots, individual CUT mutants with pink dots, the sextuple CUT mutant with purple dots, and other compound CUT mutants with green dots. One-way ANOVA followed by Tukey’s multiple comparisons test; *P < 0.05, **P < 0.01, ***P < 0.001. n ≥ 10 for all genotypes. All genotypes were compared, but only those comparison that show statistically significant differences are indicated with lines. **(D-E)** Expression of *ric-4(syb2878[ric-4::GFP])* (*ric-4(syb2878[ric-4::T2A-3xNLS-GFP])*) (D), *egl-3(syb4478[egl-3::GFP])* (*egl-3(syb4478[egl-3::SL2-GFP-H2B])*) (E) in wild-type (top) and CUT sextuple mutant (bottom). Lateral views of the worm head at the L4 stage are shown. Quantification of CRISPR alleles fluorescence intensity in head neurons. The data are presented as individual values with each dot representing the expression level of one worm with the mean ± SEM indicated. Unpaired *t*-test, ***P < 0.001. n ≥ 12 for all genotypes. **(F)** Expression of *rab-3prom::2xNLS-tagRFP[otIs356]* was compared between wild-type, CUT sextuple mutant, and CUT sextuple mutant rescue (pan-neuronal, *ceh-48* promoter (“neu”), or ubiquitous, *eft-3* promoter (“ubi”), expression of *ceh-48*, *ceh-44*, *ceh-38*, *ceh- 39* or *hOC1*). Quantification of fluorescence intensity analyzed by COPAS system (“worm sorter”). The data are presented as individual values with each dot representing the expression level of one worm with the mean ± SEM indicated. Wild-type data is represented with black dots, the sextuple CUT mutant with purple dots, and rescue lines with blue dots. One-way ANOVA followed by Tukey’s multiple comparisons test; ***P < 0.001. n ≥ 40 for all genotypes. **(G-J)** Neurotransmitter reporter transgenes in CUT gene mutants. Transgenes are *otIs518* (*eat-4fosmid::SL2::mCherry::H2B*) (G) and *otIs794* which contains *cho- 1fosmid::NLS-SL2-YFP-H2B* (H), *unc-47prom::tagBFP2* (I), and *cat-1prom::mMaroon* (J), analyzed in a wild-type (left) or CUT sextuple mutant background (right). Lateral views of the worm head at the L4 stage are shown. Quantification of fluorescence intensity in head neurons. The data are presented as individual values with each dot representing the expression level of one worm with the mean ± SEM indicated. n ≥ 10 for all genotypes. WT, wild-type; a.u., arbitrary units. Scale bars 15 μm.

ChIP analysis from the modENCODE project shows that the conserved and ubiquitously expressed CEH-38 ONECUT protein displays the same binding profile to pan-neuronal genes as the CEH-48 protein (Davis et al., 2018) (**Table S1**). Moreover, motif extraction from the ChIP-seq data reveals that CEH-48 and CEH-38 consensus binding motifs are identical (**Figure 2A**). To test the possibility that CEH-38 could compensate for loss of *ceh-44* and *ceh-48*, we generated a triple *ceh-44*; *ceh-48*; *ceh-38* null mutant strain and indeed now found a reduction of *rab-3/RAB3* expression (**Figure 3A**). Since *rab-3/RAB3* expression was not entirely eliminated, and since the *ceh-38* result indicates that even a ubiquitously expressed CUT gene contributes to the regulation of pan-neuronal gene expression, we also considered a role of the three remaining, ubiquitously expressed CUT genes that are located in an operon (*ceh-41*, *ceh-21* and *ceh-39*; **Figure 1D**). We used CRISPR/Cas9 to generate a precise deletion of those three genes and found that this deletion (*otDf1*; **Figure 1D**) alone has no significant effect on *rab-3* reporter expression (**Figure 3A**). However, adding this deletion into the context of the *ceh-44; ceh-48; ceh-38* triple mutant revealed that the sextuple CUT mutant strain displayed the strongest effect on *rab-3* expression throughout the nervous system (**Figure 3A**). Sextuple CUT mutants further displayed a significant reduction in the expression of four other pan-neuronal genes, *unc-11/SNAP91*, *ric-19/ICA1, ric-4/SNAP25* and *egl-3/PCSK2* (**Figure 3B-3E**). We tested two of these additional pan- neuronal genes, *unc-11/SNAP91* and *ric-19/ICA1,* for whether they show cumulative expression defects upon removal of individual and multiple CUT genes in combination and found this to be the case (**Figure 3B and 3C**). The joint involvement of multiple CUT genes provides an explanation for why previous screens for mutants affecting pan- neuronal gene expression were unsuccessful (Leyva-Diaz et al., 2017) and are a testament to the robustness of pan-neuronal gene expression control.

Defects observed in the compound CUT mutants are complementary to the gene expression defects observed in neuron type-specific terminal selector mutants. Specifically, genes that are more selectively expressed in the nervous system, including *cho-1/ChT* (a marker that is exclusive to cholinergic neurons), *eat-4/VGLUT* (a marker specific to glutamatergic neurons), *unc-47/VGAT* (a marker specific for GABAergic neurons) and *cat-1/VMAT* (monoaminergic neuron marker), were not affected in sextuple CUT mutant animals (**Figure 3G-3J**). This result is consistent with these genes lacking ChIP peaks of CUT protein binding (**Table S1**). Thus, the sextuple CUT mutant phenotype is a mirror image of the phenotype of terminal selector transcription factors, whose removal results in loss of neuron type-specific identity features (such as the tested *cho- 1/ChT, eat-4/VGLUT, unc-47/VGAT, cat-1/VMAT)*, but not pan-neuronal identity features (Hobert, 2016).

As expected from a loss of pan-neuronal gene expression, sextuple CUT mutant animals are severely deficient in nervous system function (**Figures 4A-B and E**). Animals display an almost complete paralysis in swimming assays, a very sensitive and well quantifiable read-out of animal locomotion (**Figure 4A**)(Croll, 1975; Pierce-Shimomura et al., 2008; Restif et al., 2014). Crawling behavior on an agar surface, quantified using a semi-automated WormTracker system, is also severely affected in sextuple CUT mutant animals (**Figure 4B**). Synaptic transmission defects, scored via responsiveness to aldicarb, are also very obvious, CUT sextuple mutants display a strong resistance to aldicarb (**Figure 4E**). We have found that these crawling and synaptic transmission defects are again cumulative, i.e. worsen the more CUT genes are removed (**Figure 4B and E**). Overall nervous system anatomy is unaffected in CUT sextuple mutants, including general cell body and fascicle position, (**Figure S4**). However, defects can be observed in synapse abundance in CUT gene mutants (**Figure 4H-I**).

**Figure 4.**
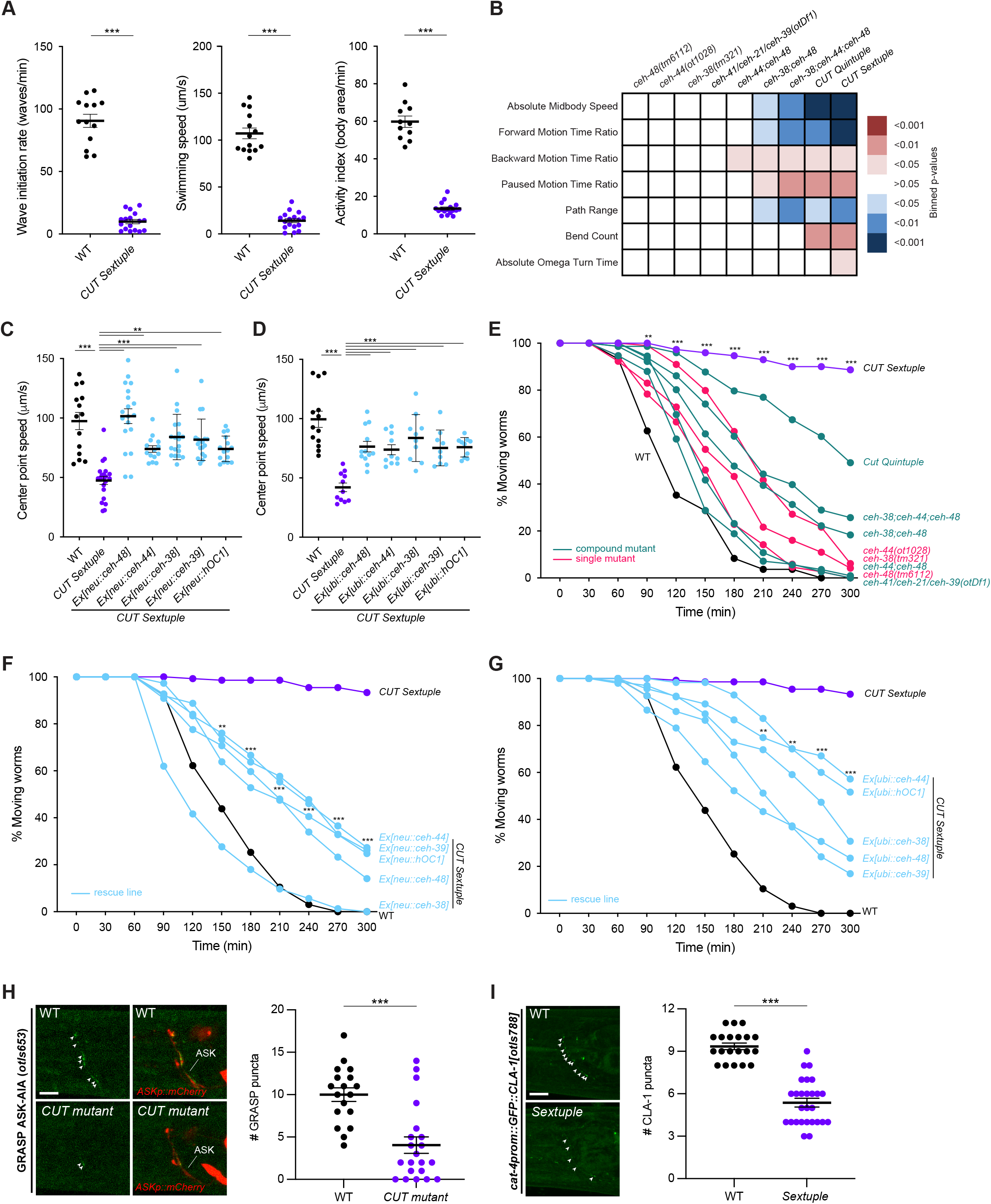
CUT genes are required for proper neuronal function. **(A)** Swimming behavior: wave initiation rate (left), swimming speed (center), and activity index (right) were compared between wild-type and CUT sextuple mutant using a multi- worm tracker system (Roussel et al., 2014). The data are presented as individual values with each dot representing the value of one worm with the mean ± SEM indicated. Unpaired *t*-test, ***P < 0.001. n ≥ 11 for all genotypes. **(B)** Behavioral phenotypic summaries of representative locomotion features for individual and compound CUT mutants, analyzed using an automated worm tracker system (Yemini et al., 2013). Heat map colors indicate the p-value for each feature for the comparison between each of the mutant strains and the wild-type strain. Red indicates a significant increase for the tested feature, while blue indicates a significant decrease. One-way ANOVA followed by Tukey’s multiple comparisons test. n ≥ 10 for all genotypes. Time ratio = (total time spent performing behavior)/(total assay time). **(C-D)** Worm speed was compared between wild-type, CUT sextuple mutant, and CUT sextuple mutant rescue (panneuronal, *ceh-48* promoter (“neu”) (**C**), or ubiquitous, *eft-3* promoter (“ubi”) (**D**), expression of *ceh-48*, *ceh-44*, *ceh-38*, *ceh-39* or *hOC1*) using a multi- worm tracker system (Roussel et al., 2014). The data are presented as individual values with each dot representing the speed of one worm with the mean ± SEM indicated. Wild- type data is represented with black dots, the sextuple CUT mutant with purple dots, and rescue lines with blue dots. One-way ANOVA followed by Tukey’s multiple comparisons test, comparisons with CUT sextuple mutant indicated; **P < 0.01, ***P < 0.001. n ≥ 10 for all genotypes. **(E)** Aldicarb-sensitivity defects in individual CUT mutants (*ceh-48(tm6112)*, *ceh-44(ot1028)*, *ceh-38(tm321)*) and compound CUT mutants (*otDf1*, which deletes *ceh-41*, *ceh-21* and *ceh-39*; double *ceh-44;ceh-48*, double *ceh-38;ceh-48*, triple *ceh-38;ceh-44;ceh-48*, quintuple *ceh-38;ceh-48;otDf1*, and sextuple *ceh-38;ceh-44;ceh-48;otDf1*) compared to wild-type animals. Aldicarb is an acetylcholinesterase inhibitor that paralyzes worms. Decreased sensitivity to aldicarb correlates with a reduction in synaptic transmission (Mahoney et al., 2006). Worms were tested every 30 min for paralysis by touching the head and tail three times each. The data are presented as the percentage of moving worms at the indicated time point, dots represent the mean of independent experiments for each genotype. Wild-type data is represented with black dots, individual CUT mutants with pink dots, the sextuple CUT mutant with purple dots, and other compound CUT mutants with green dots. Two-way ANOVA followed by Tukey’s multiple comparisons test, comparisons for wild-type vs CUT sextuple mutant indicated; **P < 0.01, ***P < 0.001. n ≥ 3 independent experiments (25 animals per independent experiment). **(F-G)** Aldicarb-sensitivity defects in wild-type animals, CUT sextuple mutant, and CUTz sextuple mutant rescue lines (pan-neuronal (F), or ubiquitous (G) rescue lines). Wild-type data is represented with black dots, the sextuple CUT mutant with purple dots, and rescue lines with blue dots. Two-way ANOVA followed by Tukey’s multiple comparisons test, comparisons for CUT sextuple mutant vs *Ex[neu::ceh-44]* (F), and CUT sextuple mutant vs *Ex[ubi::ceh-44]* (G) indicated; **P < 0.01, ***P < 0.001. n ≥ 3 independent experiments (25 animals per independent experiment). **(H)** ASK-AIA GRASP signal for the ASK>AIA (*otIs653*) in wild-type (top) and CUT compound mutant (*ceh-38(tm321)*; *ceh-44(ot1028); otDf1*) (bottom). Lateral views of L1 worm heads at the nerve ring level are shown. ASK axon is labelled with cytoplasmic mCherry. Arrowheads indicate GRASP GFP synaptic puncta. *otIs653* and *ceh-48* are located in the same chromosome (chr. IV) and cannot be recombined together. Quantification of puncta along the ASK axon in the nerve ring. The data are presented as individual values with each dot representing the number of puncta in one worm with the mean ± SEM indicated. Unpaired *t*-test, ***P < 0.001. n ≥ 18 for all genotypes. **(I)** HSN presynaptic specializations labeled by GFP-CLA-1 (*cat-4prom::GFP::CLA- 1[otIs788]*) in wild-type (top) and CUT sextuple mutant (bottom). Lateral views of young adult worm heads at the nerve ring level are shown. Arrowheads indicate CLA-1 presynaptic specializations. Quantification of CLA-1 puncta along the HSN axon in the nerve ring. The data are presented as individual values with each dot representing the number of puncta in one worm with the mean ± SEM indicated. Unpaired *t*-test, ***P < 0.001. n ≥ 20 for all genotypes. WT, wild-type; Scale bars 5 μm.

The cumulative effects of CUT gene removal suggest a scenario in which it is primarily the overall dosage of CUT genes, rather than specific features of each individual CUT gene that is important to specify pan-neuronal gene expression. To further test this notion, we re-introduced individual CUT genes into the sextuple CUT mutant background. We used two separate drivers – a ubiquitous driver (*eft-3prom*) or a pan-neuronal driver (*ceh-48prom4*, **Figure S1B**) – to generate multicopy transgenic arrays for overexpression. We found that each individually tested, overexpressed *C. elegans* CUT gene is alone able to rescue (a) the pan-neuronal gene expression defects (**Figure 3F**) and (b) the crawling and synaptic transmission defects of sextuple mutant animals (**Figures 4C-D, 4F-G and S3A-D**). Moreover, overexpression of a human ONECUT homolog, *hOC1*, is also capable of rescueing the *C. elegans* CUT sextuple mutant phenotype, indicating a conservation of gene function (**Figures 3F, 4C-D, 4F-G and S3A-D**). These results permit three conclusions: First, the usage of the postmitotic, pan- neuronal *ceh-48* promoter indicates that CUT genes indeed act cell-autonomously in postmitotic neurons; second, CUT genes are functionally interchangeable; and, third, CUT gene dosage in the nervous system appears to be the main determinant of CUT gene function as regulators of pan-neuronal gene expression.

### Raising CUT gene dosage in non-neuronal cells results in induction of pan- neuronal gene expression

We asked whether raising CUT gene dosage in non-neuronal cells is sufficient to induce pan-neuronal gene expression in non-neuronal cells. We had difficulties in establishing transgenic lines that overexpress CUT genes in non-neuronal cells using a ubiquitous driver (from the *eft-3* gene) and therefore refrained to raising CUT gene dosage in a non-essential cell type that is easy to target. To this end we made use of the promoter of the *mir-228* gene, whose expression is largely restricted to glia cells (Pierce et al., 2008). We find that overexpression of *ceh-48* in glia cells results in the ectopic induction of the pan-neuronal *rab-3* gene (**Figure 5**). This result suggests that ectopic expression of neuronal CUT genes, increasing overall CUT factors availability, might be sufficient to induce pan-neuronal features in other non-neuronal cells.

**Figure 5.**
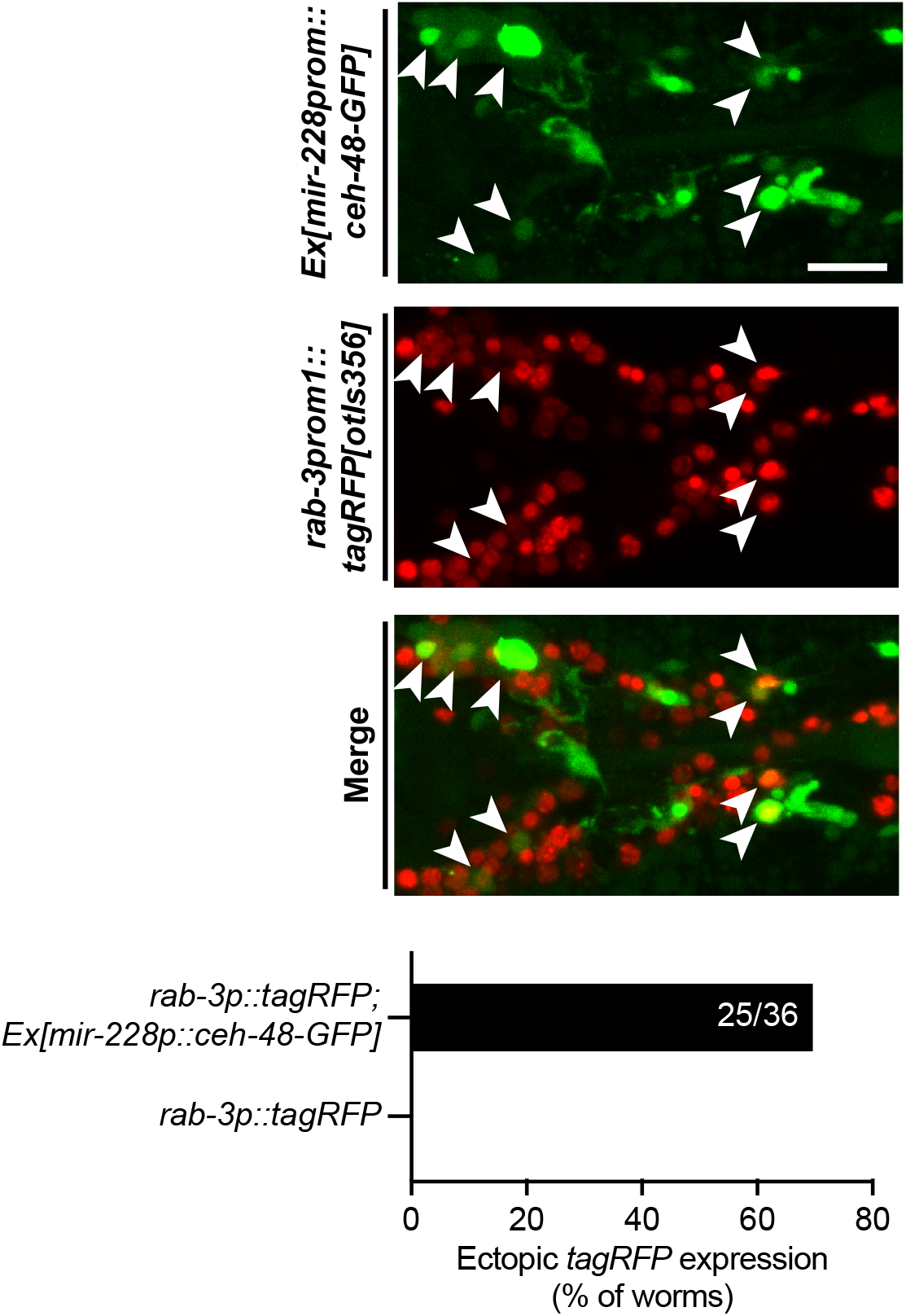
Ectopic expression of *ceh-48* induces pan-neuronal features in glia cells. Expression of *ceh-48* in glia cells under a glia specific promoter (top, *mir-228prom::ceh- 48::GFP[otEx7481]*) results in ectopic expression of the pan-neuronal marker *rab-3* (middle, *rab-3prom1::2xNLS-tagRFP[otIs356]*). The overlap (bottom) reveals multiple glia cells expressing *rab-3* in the worm head (arrowheads). Graph represents the percentage of *mir-228prom::ceh-48::GFP[otEx7481]* worms with ectopic *rab-3prom1::2xNLS- tagRFP[otIs356]* expression in glia cells. Absence of ectopic *rab-3* expression in *rab- 3prom1::2xNLS-tagRFP[otIs356]* control animals was confirmed using a pan-glial *mir- 228prom::GFP[nsIs198]* marker. Scale bar 5 μm.

### Genome-wide analysis of Cut homeobox gene targets

We further expanded our characterization of CUT gene function by RNA transcriptome profiling of CUT gene mutant animals. To this end, we used Isolation of Nuclei TAgged in specific Cell Types (INTACT) technology (Steiner et al., 2012; Sun and Hobert, 2021) to isolate all neuronal nuclei and compared neuronal transcriptomes of wild-type animals with those of sextuple CUT mutant animals (**Figure 6A)**. Apart from upregulated genes, we found >2,000 genes to be downregulated (FDR < 0.05) and about 605 (29%) of those have CUT binding ChIP peaks (**Figure 6B and 6C, Table S2**). Downregulated genes with CUT binding peak include known pan-neuronally expressed genes involved in the synaptic vesicle cycle (e.g. *unc-57/SH3GL3, ric-19/ICA1, unc- 11/SNAP91*), synaptic activity zone assembly (e.g. *cla-1/PCLO*), neuronal transport (e.g. *unc-116/JIP3*), axon pathfinding (e.g. *unc-14/RUSC1*), neuronal cytoskeleton (*e.g. unc- 119/UNC119, unc-69/SCOC*), neuropeptide processing (*e.g. egl-3/PCSK2, egl-21/CPE, pamn-1/PAM*) and other previously known pan-neuronal genes (e.g. *rgef-1/RASGRP3*, a commonly used pan-neuronal marker). Focusing on the battery of 23 pan-neuronal genes whose expression patterns we had defined in a previous analysis (Stefanakis et al., 2015), we found that most of them show reduced transcript levels in the CUT sextuple mutant (**Figure 6D**). As described above, we have validated these changes in expression for *rab-3*, *unc-11*, *ric-19*, *ric-4* and *ric-19* (**Figure 3A-E**).

**Figure 6.**
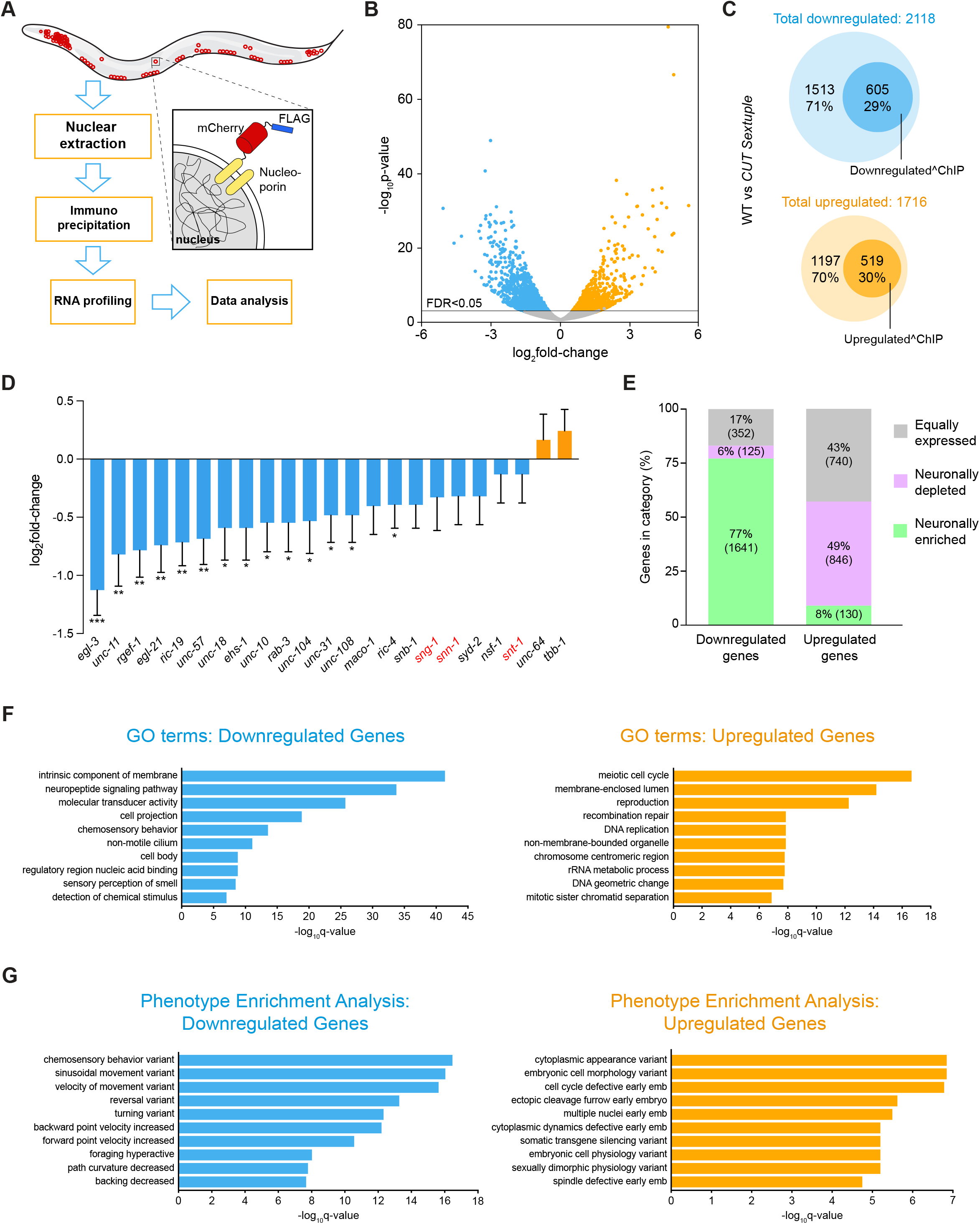
Transcriptional profiling of CUT sextuple mutants. **(A)** Schematic and experimental design for INTACT sample collection, protocol, and data analysis for neuronal transcriptome profiling. **(B)** Volcano plot of differentially expressed genes in CUT sextuple mutant neurons showing significantly (FDR < 0.05) downregulated (blue) or upregulated (orange) genes (RNA-seq, n = 3). See **Table S2** for full list of differentially expressed genes. **(C)** Diagrams showing overlap between differentially expressed genes in CUT sextuple mutant and genes bound by CEH-48 or CEH-38 in a wild-type ChIP-seq (Davis et al., 2018). Downregulated genes are marked in blue, upregulated genes are marked in orange, and the genes that contain CUT peaks are marked with dark circles within both clusters. **(D)** Changes of previously described pan-neuronal gene battery (Stefanakis et al., 2015) in CUT sextuple mutant animals. The data are presented as the log2FoldChange ± standard error calculated by DESeq2, comparing neuronal samples from wild-type and CUT sextuple mutant. The two-stage step-up method of Benjamini, Krieger and Yekutieli (FDR 10%) was used to calculate the q-values for this subset of genes, analyzing the individual p-values obtained from the DESeq2 comparison. *Q < 0.05, **Q < 0.01, ***Q < 0.1 (RNA-seq, n = 3). **(E)** Vertical slices representation of the distribution (in percentage) of the downregulated and the upregulated gene sets between the neuronally enriched (green), neuronally depleted (purple) and equally expressed (gray) gene sets. See **Table S3** for full list of neuronally enriched and depleted genes. **(F-G)** GO enrichment analysis (A) and phenotype enrichment analysis (B) using gene sets of significantly downregulated (blue) or upregulated (orange) transcripts. Graphs illustrate the 10 most significant terms. Analysis performed using the Gene Set Enrichment Analysis Tool from Wormbase. See **Table S4** for full list of enriched terms. WT, wild-type.

The use of INTACT technology to isolate the entire nervous system from wild-type animals allowed us to identify 6372 neuronally enriched genes through comparison of neuronal-nuclei to total nuclei samples (**Figure 6E, Table S3**). Among the differentially expressed genes in CUT sextuple mutants, a large proportion (77%) of the downregulated gene set corresponds to this neuronally enriched gene set, while only 8% of the upregulated genes belong to the neuronally enriched gene set. Around half of the upregulated genes are actually neuronally depleted genes, whereas the other half corresponds to genes equally distributed between the nervous system and the whole animal (**Figure 6E, Table S3**). Moreover, the downregulated gene set, but not the upregulated set, display a significantly GO term enrichment for several neuronal processes (e.g. neuropeptide signaling pathway, chemosensory behavior)(**Figure 6F; Table S4)**. Similarly, phenotype enrichment analysis for the downregulated, but not upregulated gene set shows a large amount of locomotion phenotypes (**Figure 6G; Table S4**). These findings are consistent with our reporter gene analysis, as well as our behavioral analysis, confirming that CUT homeodomain proteins are critical activators of pan-neuronal genes essential for proper neuronal function.

We find that the expression of some ubiquitously expressed genes, with potential selective functions in the nervous system, can also be CUT gene dependent. For example, we find that the *C. elegans* orthologs of the vertebrate neuronal splicing regulator *NOVA1* (Jensen et al., 2000), the *C. elegans* ortholog of the alternative splicing factor *RBM25*, and the *C. elegans* homolog of a regulator of endocytosis, *EPS15* (Ioannou and Marat, 2012), show diminished transcript levels in the transcriptome analysis of CUT sextuple mutants. All three loci show binding of CUT proteins by ChIP analysis in the modENCODE dataset (**Table S1**). *gfp* reporter alleles that we generated using CRISPR/Cas9-genome engineering revealed ubiquitous expression of *nova-1, rbm-25* and *ehs-1* throughout all tissue types (**Figure S5A-S5C**). We confirmed the CUT dependence of these genes in two different manners. First, we crossed the *nova-1* reporter allele into a CUT sextuple mutant background and found diminished expression in the nervous system. Second, we deleted the CUT binding site from the ubiquitously expressed *ehs-1* gene locus and also observed diminished expression in the nervous system (**Figure S5D**). Intriguingly, in the case of *ehs-1* this downregulation was specific to the nervous system since non-neuronal cells did not show downregulation (**Figure S5E**). Taken together, these results demonstrate the critical role of CUT-dependent gene expression of even ubiquitously expressed genes.

### Collaboration of Cut homeobox genes with terminal selectors

One notable feature of our CUT gene mutant analysis is that even in the sextuple CUT mutant, pan-neuronal gene expression is not uniformly eliminated. Nor do sextuple mutants display the larval lethality observed upon genetic removal of synaptic transmission machinery (Kohn et al., 2000). To address this conundrum, we considered our previous functional analysis of neuron-type specific terminal selectors, which are required for the initiation of neuron-type specific gene expression profiles (Hobert, 2016; Hobert et al., 2010; Hobert and Kratsios, 2019; Stefanakis et al., 2015). While terminal selector removal alone does not generally affect pan-neuronal gene expression, we had found that pan-neuronal genes do contain terminal selector binding sites, and we had shown that these binding sites are functionally relevant, but only in the context of isolated *cis*-regulatory elements (Stefanakis et al., 2015). Based on these findings, we had suggested that terminal selectors may provide redundant regulatory input into pan- neuronal gene expression (**Figure 7A**)(Stefanakis et al., 2015). Hence, an explanation for the lack of a complete loss of pan-neuronal gene expression in CUT sextuple mutants would be that terminal selectors are responsible for residual pan-neuronal gene expression.

**Figure 7.**
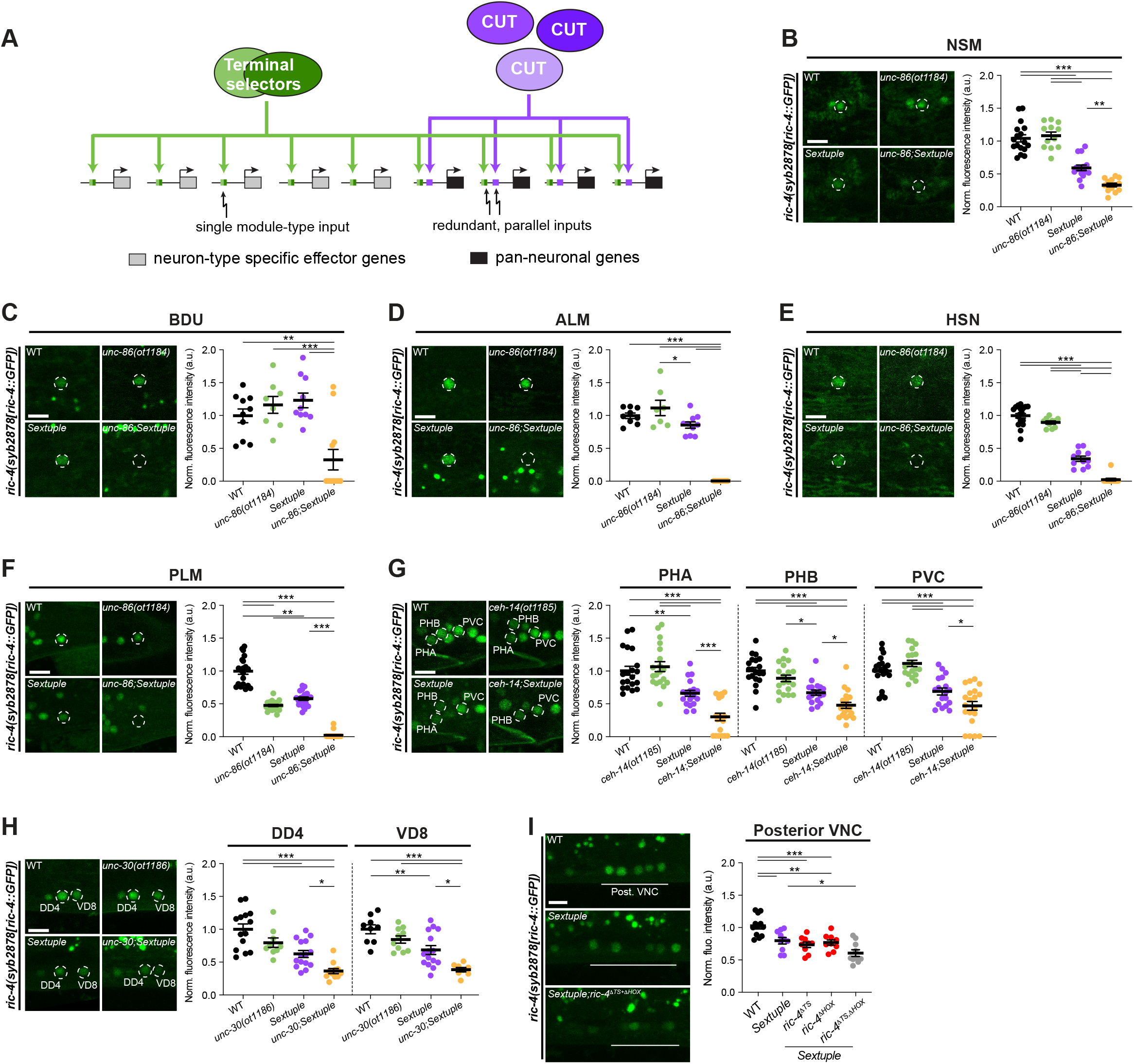
CUT genes cooperate with terminal selectors to control pan-neuronal gene expression. **(A)** Illustration for how terminal selectors contribute to the regulation of pan-neuronal gene expression. **(B-H)** Expression of *ric-4(syb2878[ric-4::GFP])* (*ric-4(syb2878[ric-4::T2A-3xNLS-GFP])*) in wild-type (top left), terminal selector mutant (*unc-86(ot1184)* (B-F), *ceh-14(ot1185)* (G) or *unc-30(ot1186)* (H); top right), CUT sextuple mutant (bottom left), and compound terminal selector and CUT sextuple mutant (bottom right). Lateral views of the head (B), midbody (C-E, F) and tail (F and G) are shown. All images correspond to worms at the L4 larval stage, except for HSN (E) where young adults are shown. Quantification of *ric- 4(syb2878[ric-4::T2A-3xNLS-GFP])* fluorescence intensity in individual neurons. The data are presented as individual values with each dot representing the expression level of NSM (B), BDU (C), ALM (D), HSN (E), PLM (F), PHA, PHB, PVC (G), DD4, or VD8 (H) neuron, with the mean ± SEM indicated. Wild-type data is represented with black dots, terminal selector mutants with green dots, the sextuple CUT mutant with purple dots, and compound terminal selector and CUT sextuple mutant with yellow dots. One-way ANOVA followed by Tukey’s multiple comparisons test; *P < 0.05, **P < 0.01, ***P < 0.001. n ≥ 8 for all genotypes. **(I)** Expression of *ric-4(syb2878[ric-4::T2A-3xNLS-GFP])* in wild-type (top), CUT sextuple mutant (middle), and upon mutation of HOX and terminal selector binding sites on the *ric- 4* endogenous locus in a CUT sextuple mutant background (bottom). Individual mutation of the HOX (*ric-4(ot1182 syb2878)*) or terminal selector binding sites (*ric-4(ot1181 syb2878)*) has no effect on *ric-4(syb2878[ric-4::T2A-3xNLS-GFP])* expression, but the expression is reduced in posterior ventral nerve cord (VNC) neurons when binding site mutations are combined (*ric-4(ot1183 ot1181 syb2878)*). Lateral views of the posterior VNC in L4 worms are shown. Quantification of *ric-4(syb2878[ric-4::T2A-3xNLS-GFP])* fluorescence intensity in posterior VNC neurons. The data are presented as individual values with each dot representing the expression level of one worm with the mean ± SEM indicated. Wild-type data is represented with black dots, the sextuple CUT mutant with purple dots, the sextuple mutant with individual binding sites mutated with red dots, and the sextuple mutant with both binding sites mutated with gray dots. One-way ANOVA followed by Tukey’s multiple comparisons test; *P < 0.05, **P < 0.01, ***P < 0.001. n ≥ 8 for all genotypes. WT, wild-type; a.u., arbitrary units. Scale bars 5 μm.

We addressed this possibility by generating different septuple null mutant strains in which we jointly removed all six CUT genes together with different terminal selectors that were previously found to regulate distinct neuron type-specific gene batteries. We indeed found that joint removal of terminal selectors and CUT genes strongly enhanced the reduction of pan-neuronal gene expression. For example, pan-neuronal gene expression in CUT sextuple mutants in ALM/PLM, HSN, BDU and NSM is further reduced, if not completely eliminated, upon removal of the POU homeobox gene *unc-86*, which is a terminal selector of these neuron classes (Leyva-Diaz et al., 2020) (**Figure 7B- 7F**). Similarly, the CUT sextuple effect in the PVC, PHA and PHB tail neurons is enhanced upon genetic removal of the LIM homeobox gene *ceh-14*, the terminal selector of PVC, PHA and PHB (**Figure 7G**)(Kagoshima et al., 2013; Pereira et al., 2015; Serrano-Saiz et al., 2013). Likewise, the DD and VD GABAergic motor neurons of the ventral nerve cord, which lose neuron-type specific identity features, but not pan-neuronal identity features upon removal of the *unc-30* Pitx homeobox gene (Cinar et al., 2005; Yu et al., 2017), show a further reduction of pan-neuronal gene expression in a septuple CUT; *unc-30* mutant background, compared to the CUT sextuple or *unc-30* single mutant background alone (**Figure 7H**).

As an independent approach to removal of a terminal selector-encoding locus, we also mutated terminal selector binding sites in a pan-neuronal gene locus and asked whether this would enhance the effect of removal of CUT genes. Indeed, mutating binding sites for terminal selectors for ventral nerve cord motor neurons into a *gfp-*tagged *ric- 4/SNAP25* locus, further decreased *ric-4/SNAP25* expression in a CUT sextuple null mutant background. (**Figure 7I)**. These results indicate that CUT factors act in concert with terminal selectors to control pan-neuronal gene batteries.

## DISCUSSION

We have shown here how a critical, but previously little understood component of neuronal gene expression programs – the expression of pan-neuronal gene batteries - is controlled. We identified an entire family of transcription factors, the CUT homeodomain transcription factors, as key regulators of pan-neuronal gene expression. CUT homeobox genes are also candidate regulators of pan-neuronal gene expression in other organisms. *Drosophila,* sea urchin and the simple chordate *Ciona intestinalis* contain a single ONECUT gene with strikingly restricted, pan-neuronal gene expression (Lowe and Stolfi, 2018; Nguyen et al., 2000; Poustka et al., 2004). In vertebrates, CUX and ONECUT gene numbers have expanded and display complex expression patterns within and outside the nervous system (Kropp and Gannon, 2016; Weiss and Nieto, 2019). A truly nervous system-wide analysis of the function of CUX or ONECUT genes in vertebrates has been missing; instead, several of vertebrate CUT or ONECUT homeodomain proteins have been well-studied in specific cellular contexts (Kropp and Gannon, 2016; Weiss and Nieto, 2019). Encouragingly, a recent analysis of *Ciona* ONECUT revealed changes in gene expression of synaptic transmission molecules upon manipulation of ONECUT function in photoreceptor differentiation (Vassalli et al., 2021). Our genetic loss of function analysis in *C. elegans* predicts that compound mutants may need to be generated in mice to assess CUT family function in vertebrate pan-neuronal gene expression.

The identification of CUT genes as regulators of pan-neuronal genes in *C. elegans* provides a complement to the much better understood regulation of neuron type-specific gene batteries. Pan-neuronal genes require at least two distinct sets of direct regulatory inputs to initiate (and presumably also maintain) their expression: a proper dosage of broadly expressed CUT homeobox genes and neuron type-specific terminal selector transcription factors (**Figure 7A**). Only the cumulative removal of all these regulatory inputs results in strong disruptions of pan-neuronal gene expression, illustrating a striking robustness of pan-neuronal gene regulation. The robustness of pan-neuronal gene regulatory architecture contrasts with the regulation of neuron-type specific gene batteries, where removal of individual *cis-*regulatory elements, or individual terminal selector transcription factors that act through such *cis-*regulatory elements, completely eliminates expression of neuron type-specific genes (Hobert, 2016; Stefanakis et al., 2015). These dichotomous regulatory strategies may speak to (a) the evolvability of neuron type-specific gene expression programs and (b) evolutionary stability of pan- neuronal gene batteries. Brain evolution involves an increase in neuronal cell type diversity, and is essentially a “variation on a theme” process, characterized by an increase in neuronal cell type diversity, in which certain parameters remain stable (pan-neuronal identity), while others rapidly evolve. The two distinct regulatory strategies for neuron type-specific and pan-neuronal gene expression may lie at the basis of such evolutionary plasticity and stability.

Our studies underscore the centrality of homeobox genes in controlling multiple aspects of neuronal identity, not just in terms of conferring neuron-type specific features as has been shown before (Hobert, 2021; Reilly et al., 2020), but also in broadly defining what distinguishes non-neuronal from neuronal cells, a cell type that has gained the ability to communicate with others via a shared synaptic machinery and neuropeptides. These points indicate that the homeobox gene family may have been recruited into the control of neuronal gene expression very early in the evolution of nervous systems.

## ACKNOWLEDGEMENTS

We thank Chi Chen for generating transgenic lines, HaoSheng Sun for help with the INTACT protocol, Maryam Majeed for providing the HSN CLA-1 and GRASP ASK-AIA strains, and for help with the analysis of GRASP data, Eviatar Yemini for help with worm tracking, Cyril Cros and Molly Reilly for providing the neurotransmitter strain, Berta Vidal for providing the *rab-3* cytoplasmic strain, Waleed Ali, Kevin Gonzalez and Mayeesa Rahman for genotyping and strain maintenance, Hynek Wichterle, Wes Gruber, Esteban Mazzoni and members of the Hobert lab for comments on this manuscript, and Wormbase and CGC (funded by NIH Office of Research Infrastructure Programs, P40 OD010440) for providing resources and reagents. This work was funded by the Howard Hughes Medical Institute. E.L.-D. was supported by an EMBO long-term fellowship.

## AUTHOR CONTRIBUTIONS

E.L-D. and O.H. conceived the project and designed the experiments. E.L-D. performed the experiments, imaging and quantifications. The manuscript was prepared by E.L-D. and O.H.

## STAR METHODS

### RESOURCE AVAILABILITY

#### Lead Contact

Further information and requests for resources and reagents should be directed to and will be fulfilled by the Lead Contact, Oliver Hobert.

#### Materials Availability

All newly generated strains will be available at the *Caenorhabditis* Genetics Center (CGC).

#### Data and Code Availability

Raw and processed RNA-seq data will be available at GEO accession #GSE188489.

### EXPERIMENTAL MODEL AND SUBJECT DETAILS

#### Caenorhabditis elegans strains and handling

Worms were grown at 20°C on nematode growth media (NGM) plates seeded with *E. coli* (OP50) bacteria as a food source unless otherwise mentioned. Worms were maintained according to standard protocol (Brenner, 1974). Wild-type strain used is Bristol variety, strain N2. A complete list of strains and transgenes used in this study is listed in the Key Resources Table.

### METHOD DETAILS

#### C. elegans strains

Except strains described below, all strains were previously published, and/or obtained from CGC or the National BioResource Project (NBRP, Japan), and/or crosses with these strains as detailed in the Key Resources Table.

#### CRISPR/Cas9-based genome engineering

*ceh-48(ot1125[ceh-48::GFP])*, *ceh-44(ot1028)*, *otDf1*, *rab-3(ot1178 syb3072)*, *unc- 10(ot1180 syb2898 syb3252)*, *ric-4(ot1123 syb2878)*, *ric-4(ot1179 ot1123 syb2878)*, *ric- 4(ot1181 syb2878)*, *ric-4(ot1182 syb2878)*, *ric-4(ot1183 ot1181 syb2878)*, *unc- 86(ot1184)*, *ceh-14(ot1185)*, *unc-30(ot1186)* were generated using Cas9 protein, tracrRNA, and crRNAs from IDT, as previously described (Dokshin et al., 2018). For *ceh-48(ot1125[ceh-48::GFP])*, one crRNA (atatgattattaggtgatta) and an assymetric double stranded *GFP-loxP-3xFLAG* cassette, amplified from a plasmid, were used to insert the fluorescent tag at the C-terminal. For *ceh-44(ot1028)*, two crRNAs (ttaaggcgacgaagttatga and ccgaggaggcgaacagctat) and a ssODN donor (ataatatgatttctataattaaggcgacgaagttatatcggcagaagaatacggattctgaacttattga) were used to delete 80 bp of *ceh-44* exon 8, introducing a frameshift in the CUT isoform (isoform a). For *otDf1*, two crRNAs (ggcatacatcttttcgaaag and atgaagaaaattatcaggat) and a ssODN donor (gaaaagggaattcggaaatgaagaaaattatcagtcgaaaagatgtatgcccgaaatgttccgagaaac) were used to generate a 8968 bp deletion (from position -159 upstream *ceh-39* ATG, to 89 bp downstream *ceh-41* stop codon) affecting 4 genes (deficiency, Df). The genes deleted in *otDf1* are *ceh-41*, *ceh-21*, *T26C11.9* and *ceh-39*. For *rab-3(ot1178 syb3072)*, one crRNA (gctcacaaaaatggatcgat) and a ssODN donor (ctatctctctccgtgagcaacgagctagtcaacccaaaaaaccatttttgtgagcacacacagagagagactcaaa) were used to mutate a CUT binding site on *rab-3(syb3072[rab-3::T2A::3xNLS::GFP])* CRISPR reporter (details on binding site mutations on section below). For *unc- 10(ot1180 syb2898 syb3252)*, one crRNA (tcgtgcttcacggaattgtg) and a ssODN donor (gcagagagagaaaagtagtcgtgcttcacggaattgtggagagaaaaaaagagatctcaagtcagagagcgcgagcttcgtttct) were used to mutate a CUT binding site on *unc-10(syb2898 syb3252[unc- 10::T2A::3xNLS::GFP])* CRISPR reporter. For *ric-4(ot1123 syb2878)*, one crRNA (atgagagccaatcgatacgt) and a ssODN donor (acgaagtgagccagaaagggaagcccgcacccacgtaaaaaaaaactctcatagagagaaagagagtctctgttttc tct) were used to mutate a CUT binding site (“site 1”) on *ric-4(syb2878[ric- 4::T2A::3xNLS::GFP])* CRISPR reporter. For *ric-4(ot1179 ot1123 syb2878)*, two crRNA (gaaaaatggaagtcacttgg and gggaaacagagaaaagacta) and a ssODN donor (aaatttcatataatttcccatccttcccacccccactaaggcttcatagtgcaaccttataactattagt) were used to delete a 431 bp section containing 9 CUT binding sites (“site 2”) within *ric-4* intron 1, on top of *ric-4(ot1123 syb2878)*. For *ric-4(ot1181 syb2878)*, one crRNA (ttgacgataacagagaccca) and a ssODN donor (ttgttcagtctttcccaaatttttgtgcccaatctAAAAAAAAAAAAAActctgttatcgtcaaaagtgacatcttttctttcg) were used to mutate COE (UNC-3) and UNC-30 binding sites on *ric-4(syb2878[ric- 4::T2A::3xNLS::GFP])* CRISPR reporter. For *ric-4(ot1182 syb2878)* and *ric-4(ot1183 ot1181 syb2878)*, one crRNA (cgaaaagagctcagcgaaaa) and a ssODN donor (tcttcgtgccatccattcaaacaacgcttattttaaaaaaaaaaacatttttcgctgagctcttttcgtttcgtctttcttgtttc) were used to mutate a HOX binding site on *ric-4(syb2878[ric-4::T2A::3xNLS::GFP])* or *ric-4(ot1183 ot1181 syb2878)*. For *unc-86(ot1184)*, two crRNAs (caaggtccccctcttttcca and acaacatacaatgggctacc) and a ssODN donor (tctgtctcctcccagcttcaaggtccccctcttttaccttgattctttgattagtttcgttttcgtgaac) were used to delete the entire *unc-86* locus. For *ceh-14(ot1185)*, two crRNAs (tcttggcgagtgcgatgagc and tgtactgtggagtcatgtgt) and a ssODN donor (gggacacaacattttgactcttggcgagtgcgatgcatgactccacagtacatttgaactggagaaaaac) were used to delete the entire *ceh-14* locus. For *unc-30(ot1186)*, two crRNAs (taagacggtaataatccttg and gtagtaaagttgaaaaggcg) and a ssODN donor (ccgatcactgactttgcgtaagacggtaataatcccttttcaactttactactgttcaataaacaattaa) were used to delete the entire *unc-30* locus.

*rab-3(syb3072)*, *ric-4(syb2878)*, *unc-10(syb2878)*, *egl-3(syb4478)*, *ceh-38(syb4799)*, *ceh-41(syb4901)*, *nova-1(syb4373)*, *rbm-25(syb4376)*, *ehs-1(syb4426)*, *ehs-1(syb4426 syb4716)*, were generated by SUNY Biotech. *ceh-38(syb4799)* and *ceh-41(syb4901)* were generated with the exact same *GFP-loxP-3xFLAG* cassette as in *ceh- 48(ot1125[ceh-48::GFP])* for direct comparison of CUT *gfp-*tagged CRISPR alleles.

For CUT binding site mutations, we looked for CEH-48 sites centered within the region covered by CEH-48 and/or CEH-38 ChIP peaks in *rab-3*, *ric-4*, *unc-10* and *ehs-1* regulatory regions. The CEH-48 binding motif (consensus ATCGA), is cataloged in the CIS-BP (Catalog of Inferred Sequence Binding Preferences) database (http://cisbp.ccbr.utoronto.ca/)(Weirauch et al., 2014). The CEH-48 motif matches known motifs for other ONECUT and CUX proteins (see ChIP-seq section below) (**Table S5)**. Deletions of CEH-48 binding sites were done by replacement of the binding site by adenines.

In *rab-3(syb3072[rab-3::T2A::3xNLS::GFP])*, ATCGAT (+2399, +2404) was mutated to AAAAAA. This site was centered within CEH-48 (+2326, +2452) and CEH-38 (+2211, +2719) ChIP peaks.

In *unc-10(syb2898 syb3252[unc-10::T2A::3xNLS::GFP])*, ATCGAT (-4558, -4553) was mutated to AAAAAA. This site was centered within CEH-48 (-4784, -4415) and CEH-38 (-4811, -4366) ChIP peaks.

In *ric-4(syb2878[ric-4::T2A::3xNLS::GFP])*, ATCGATTGG (-3683, -3675; “site 1”) was mutated to AAAAAAAAA. This site was centered within CEH-48 (-3832, -3598) and CEH-38 (-4062, -3521) ChIP peaks.

In *ehs-1(syb4426[ehs-1::SL2-GFP-H2B])*, ATCGAT (-220, -215) was mutated to AAAAAA. This site was centered within CEH-48 (-311, -106) and CEH-38 (-373, -168) ChIP peaks.

For *ric-4*, a second set of CUT binding sites (“site 2”) was mutated within *ric-4prom25* (*cis-*regulatory element found to be broadly expressed in head neurons)(Stefanakis et al., 2015). A 431 bp section (+4947, +5378) in *ric-4* intron 1, containing 9 CUT sites, was deleted.

The HOX/EXD motif, COE (UNC-3) motif, and UNC-30 motifs on *ric-4* were mutated following prior experiments in small *cis*-regulatory elements (Stefanakis et al., 2015), but here these mutations were done on the *ric-4* CRISPR reporter allele, *ric-4(syb2878[ric- 4::T2A::3xNLS::GFP])*. The HOX motif TGAATAATTG (-1064, -1055) was mutated to AAAAAAAAAA. The COE, TCCCTTGGGT (-1349, -1340), and UNC-30, TAATCC (-1352, -1347), motifs partially overlap and were mutated together: CTAATCCCTTGGGT was mutated to AAAAAAAAAAAAAA.

In the small *cis*-regulatory element reporters (see below) mutations in the same CUT binding sites described here for *rab-3*, *ric-4* and *unc-10* were introduced in *rab- 3prom10*, *ric-4prom30* (site 1) and *unc-10prom12*.

#### Reporter transgenes

The *rab-3*, *ric-4* and *unc-10 cis*-regulatory element reporters were generated using a PCR fusion approach (Hobert, 2002). The *rab-3prom10* (+2326, +2452) (promoter fragment number continues the series generated for *cis-*regulatory analysis in (Stefanakis et al., 2015)), *ric-4prom30* (-3832, -3598) and *unc-10prom12* (-4784, -4415) promoter fragments were amplified from N2 genomic DNA and fused to *2xNLS-GFP*. These promoter fragment coordinates match those of the CEH-48 ChIP peaks in the regulatory regions of these genes. The resulting PCR fusion DNA fragments were injected as simple extrachromosomal arrays (50 ng/mL) into *pha-1(e2123)* animals, using a *pha-1* rescuing plasmid (pBX at 50 ng/μL) as co-injection marker.

Extrachromosomal array lines were selected according to standard protocol. For *rab- 3prom10*, *ric-4prom30* and *unc-10prom12* harboring the CUT site mutations, promoters were obtained as gBlocks (IDT) and fused to *2xNLS-GFP*.

To assess neurotransmitter identity, we generated a transgene that expresses multiple reporters that assess neurotransmitter usage, including: a *cho-1* fosmid reporter construct (*cho-1fosmid::NLS-SL2-YFP-H2B*; (Stefanakis et al., 2015)), to label cholinergic neurons; an *eat-4* fosmid reporter construct (*eat- 4fosmid::SL2::mCherry::H2B* (Serrano-Saiz et al., 2013), where *mCherry* was replaced with *LSSmOrange*) to label glutamatergic neurons; *unc-47prom* (coordinates -2778, -1) fused with *TagBFP2* to label GABAergic neurons, *cat-1prom* (-1599, -1) fused with *mMaroon* to label monoaminergic neurons, and *rab-3prom1* (-1462, +2921) fused with *tagRFP* to label all neurons (pan-neuronal marker). The *cho-1fosmid::NLS-SL2-YFP- H2B* (20 ng/μL), *eat-4fosmid::SL2:: LSSmOrange::H2B* (20 ng/μL), *unc- 47prom::tagBFP2* (5 ng/μL), *cat-1prom::mMaroon* (5 ng/μL) and *rab3prom1::2xNLS- tagRFP* (10 ng/ μL) constructs were injected together, and the resulting extrachromosomal array strain was integrated into the genome using standard UV irradiation methods. This was followed by 3 rounds of backcrossing to N2 to generate *otIs794*.

To generate *cat-4prom::GFP::CLA-1(S)* (pMM13), *cat-4prom8* (-629, -299; expressed in HSN; (Lloret-Fernandez et al., 2018)) was amplified from N2 genomic DNA. The PCR fragment was cloned into PK065 (kindly shared by Peri Kurshan). *cat-4prom::mCherry* (pMM11) was generated similarly and cloned into pPD95.75. The constructs pMM13 and pMM11 were injected at 5 and 30 ng/μL, respectively, with an *inx-16prom::tagRFP* co-injection marker (10 ng/μL). The resulting extrachromosomal array strain was integrated into the genome using standard UV irradiation methods.

To label the ASK-AIA synapse with GRASP (Feinberg et al., 2008), we generated *otIs653(srg-8prom::mCherry, cho-1prom::mCherry, srg-8prom::NLG-1::spGFP1-10, cho-1prom::NLG-1::spGFP11*). For this transgene, a 2kb *srg-8prom* (coordinates -2000, -1; expressed in ASK) was cloned into MVC2 (*pSM::NLG-1::spGFP1-10*) using RF cloning to generate *srg-8prom::NLG-1::spGFP1-10* (pMM14). *srg-8prom::mCherry* (pMM02) was generated by subcloning *srg-8prom* into pPD95.75. A 364bp *cho-1prom* (- 3006, -2642; expressed strongly in AIA, AIY, AIN; (Serrano-Saiz et al., 2020)) PCR fragment amplified from genomic DNA was cloned into MVC3 (*pSM::NLG-1::spGFP11*) and pPD95.75 to generate *cho-1prom::NLG-1::spGFP11* (pMM08) and *cho- 1prom::mCherry* (pMM07), respectively. The constructs were injected at a total of 90 ng/μL, transgenic lines were picked based on the mCherry cytoplasmic expression, and the resulting extrachromosomal array strain was integrated into the genome using standard UV irradiation methods.

To generate the *rab-3* cytoplasmic reporter (*rab-3prom1::GFP*), *rab-3* promoter (“prom1” (Stefanakis et al., 2015)) was cloned into pPD95.67 (plasmid containing *2xNLS-GFP*), where the *2xNLS* was removed. The resulting plasmid was injected injected as simple extrachromosomal array (50 ng/μL) into N2 animals, using *ttx-3prom::mCherry* as a co-injection marker (25 ng/μL). The resulting extrachromosomal array strain was integrated into the genome using standard UV irradiation methods. This was followed by 6 rounds of backcrossing to N2 to generate *otIs748*.

#### CUT gene rescue and misexpression

*ceh-48* promoter (prom4, -2524, -1876) was amplified from N2 genomic DNA. *eft-3* promoter (-622, -13) was amplified from pDD104 (*eft-3prom::Cre*). *ceh-48*, *ceh-38*, *ceh- 44* and *ceh-39* cDNAs were amplified from N2 cDNA. Human *ONECUT1* (*hOC1*) cDNA was obtained from Dharmacon. The promoter fragments and the cDNAs were cloned together by Gibson assembly. The constructs were injected at 7 ng/μL as complex extrachromosomal arrays in the CUT sextuple mutant strain, and extrachromosomal array lines were selected according to standard protocol.

*mir-228* promoter (prom4, -2524, -1876) was amplified from pSW25-mir228-mCherry (kindly shared by Sean Wallace, Shaham laboratory). *ceh-48* cDNA was amplified from N2 cDNA. The promoter fragment and the cDNA were cloned together by Gibson assembly. The constructs were injected at 20 ng/μL as simple extrachromosomal arrays in *otIs356(rab-3prom1::2xNLS-tagRFP)* worms, and extrachromosomal array lines were selected according to standard protocol.

#### Automated Worm tracking

Automated single worm tracking was performed using the Wormtracker 2.0 system at room temperature (Yemini et al., 2013). Young adult animals were recorded for 5 min and tracked on NGM plates with a small patch of food in the center (5 μL OP50 bacteria). Analysis of the tracking videos was performed as previously described (Yemini et al., 2013). For the tracking of the CUT rescue lines and controls, tracking was performed using the WormLab automated multi-worm tracking system (MBF Bio- science)(Roussel et al., 2014) at room temperature. In each plate, 5 young adult animals were recorded for 5 min and tracked on NGM plates with a small patch of food in the center (5 μL OP50 bacteria). Videos were segmented to extract the worm contour and skeleton for phenotypic analysis. Raw WormLab data was exported to Prism (GraphPad) for further statistical analysis. Statistical significance between each group was calculated using One-way ANOVA followed by Tukey’s multiple comparisons test.

#### Swimming analysis

The swimming assay was performed as previously described (Restif et al., 2014) using the WormLab automated multi-worm tracking system (MBF Bio-science)(Roussel et al., 2014) at room temperature. In brief, 5 young adult animals were transferred into 50 µl M9 buffer and recorded for 1 min. Multiple features of the swim behavior were then analyzed using the WormLab software. Swimming metrics are based on the metrics described in (Restif et al., 2014). WormLab data was exported to Prism (GraphPad) for further statistical analysis.

#### Aldicarb assays

Aldicarb assays were performed as previously described (Mahoney et al., 2006). Briefly, 25 young adult animals (24 h after L4 stage, blinded for genotype) were picked into freshly seeded NGM plates containing 1 mM aldicarb (ChemService). Worms were assayed for paralysis every 30 min by prodding with a platinum wire. A worm was considered paralyzed if it did not respond to prodding to the head and tail three times each at a given time point. Strains were grown and assayed at room temperature.

Statistical significance between each group was calculated in Prism (GraphPad) using Two-way ANOVA followed by Tukey’s multiple comparisons test.

#### Microscopy

Worms were anesthetized using 100mM of sodium azide and mounted on 5% agarose on glass slides. All images were acquired using a Zeiss confocal microscope (LSM880). Image reconstructions were performed using Zen software tools. Maximum intensity projections of representative images were shown. Fluorescence intensity was quantified using the ImageJ software (Schneider et al., 2012). Figures were prepared using Adobe Illustrator.

#### INTACT for purification of affinity-tagged neuronal nuclei

UPN::INTACT control worms (*otIs790*) as well as CUT sextuple mutant were grown on large plates (150mm) with enriched peptone media coated with NA22 bacteria to allow for the growth of large quantities of worms: 100,000 worms can grow from synchronized L1 stage to gravid adults on a single plate. ∼600,000 animals were collected for each replicate at the L1 larval stage after egg preparation according to standard protocol.

Animals were washed off the plate with M9, washed 3x with M9, lightly fixed with cold RNAse-free DMF for 2 minutes before washing with 1xPBS 3x. We followed the modified INTACT protocol (Sun and Hobert, 2021) to optimize pull-down of neuronal nuclei. All steps following were done in cold rooms (4 °C) to minimize RNA and protein tag degradation. The animals were homogenized mechanically using disposable tissue grinders (Fisher) in 1x hypotonic buffer (1x HB: 10 mM Tris pH 7.5, 10 mM NaCl, 10 mM KCl, 2 mM EDTA, 0.5 mM EGTA, 0.5 mM Spermidine, 0.2 mM Spermine, 0.2 mM DTT, 0.1% Triton X-100, 1x protease inhibitor). After each round of mechanical grinding (60 turns of the grinder), the grinder was washed with 1 mL 1x HB and the entire homogenate was centrifuged at 100xg for 3 min. The supernatant was collected for later nuclei extraction and the pellet was put under mechanical grinding and centrifugation for 4 additional rounds. The supernatant collected from each round were pooled, dounced in a glass dounce, and gently passed through an 18-gauge needle 20x to further break down small clumps of cells. The supernatant was then centrifuged at 100xg for 10 min to further remove debris and large clumps of cells. Nuclei was isolated from the supernatant using Optiprep (Sigma): supernatant after centrifugation was collected in a 50mL tube, added with nuclei purification buffer (1x NPB: 10 mM Tris pH 7.5, 40 mM NaCl, 90 mM KCl, 2mM EDTA, 0.5 mM EGTA, 0.5 mM Spermidine, 0.2 mM Spermine, 0.1 mM DTT, 0.1% Triton X-100, 1x protease inhibitor) to 20 mL, and layered on top of 5 mL of 100% Optiprep and 10 mL of 40% Optiprep. The layered solution was centrifuged at 5000xg for 10 min in a swinging bucket centrifuge at 4 °C. The nuclei fraction was collected at the 40/100% Optiprep interface. After removal of the top and bottom layers, leaving a small volume containing the nuclei, the process was repeated 2 additional times. After final collection of the crude nuclei fraction, the volume was added to 4 mL with 1xNPB and precleared with 10 µL of Protein-G Dynabeads and 10 µL of M270 Carboxylated beads for 30 min to 1 h (Invitrogen). The precleared nuclei extract was then removed, and 50 µL was taken out as input samples (total nuclei). The rest was incubated with 30 µL of Protein G Dynabeads and 3 µL of anti-FLAG M2 antibody (Sigma) overnight to immunoprecipitate (IP) the neuronal nuclei. The following day, the IPed neuronal nuclei/beads was washed 6-8 times with 1xNPB for 10-15 min each time. The resulting IPed neuronal nuclei/beads were resuspended in 50 uL 1xNPB and a small aliquot was used to check with DAPI staining to quality-check the procedure for the following: 1) sufficient quantities of nuclei was immunoprecipitated; 2) nuclei are intact and not broken; 3) the majority of bound nuclei are single, mCherry-labelled neuronal nuclei and minimal nuclei clumps and large tissue chunks were immunoprecipitated. Anything not satisfying these quality checks were not used for downstream processing. The resulting input and neuronal IP samples were used for isolation of total RNA using Nucleospin RNA XS kit according to manufacturer’s protocol (Takara).

#### RNA-seq and data analysis

RNA-seq libraries were prepared using the Universal RNA-seq kit (Tecan) according to manufacturer’s protocol. The libraries were sequenced on Illumina NextSeq 500 machines with 75bp single-end reads. After initial quality check, the reads were mapped to WS220 using STAR (Dobin et al., 2013) and assigned to genes using featurecounts (Liao et al., 2014). Differential gene expression analysis was conducted using DESeq2 (Love et al., 2014). 3834 genes were found to be differentially expressed in CUT sextuple mutants compared to wild-type animals (FDF < 0.05) (**Table S2**). Gene Ontology and Phenotype Enrichment Analysis were performed using the Gene Set Enrichment Analysis tool from Wormbase (https://wormbase.org) (**Table S4**).

#### ChIP-seq datasets analysis

The CEH-48 ChIP-seq dataset (Experiment: ENCSR844VCY, bigBed file containing peak information: ENCFF784CKU) was obtained from the ENCODE portal (https://www.encodeproject.org/). The CEH-38 ChIP-seq dataset (Accession # modEncode_4800, gff3 interpreted data file containing peak information for combined replicates) was obtained from the modENCODE portal (http://www.modencode.org/). To identify the genes associated with these peak regions, peak coordinates were intersected with gene promoter regions (defined as from 5kb upstream of the transcription start site to 1kb downstream), and overlapping genes were identified (**Table S1**). The consensus binding motif for CEH-48 and CEH-38 was obtained using MEME-ChIP (Machanick and Bailey, 2011), which returned similar motifs for both factors (consensus AATCGATA). Comparison of these motifs, and of the CEH-48 motif defined in (Weirauch et al., 2014), to known motifs using the Tomtom Motif Comparison Tool in MEME Suite (Gupta et al., 2007) returned matches to known motifs for other ONECUT and CUX proteins (**Table S5**).

### QUANTIFICATION AND STATISTICAL ANALYSIS

All microscopy fluorescence quantifications were done in the ImageJ software (Schneider et al., 2012). For all images used for fluorescence intensity quantification, the acquisition parameters were maintained constant among all samples (same pixel size, laser intensity), with control and experimental conditions imaged in the same imaging session. For quantification of head neurons (**Figure 2**, **Figure 3**), nerve ring neurons (**Figure S1**, **Figure S5**) and ventral nerve cord neurons (**Figure 7I**), fluorescence intensity was measured in maximum intensity projections using a single rectangular region of interest. A common standard threshold was assigned to all the control and experimental conditions being compared. For quantification of individual neurons (**Figure 7**), fluorescence intensity was measured in the focal plane with the strongest neuronal nucleus signal within the z-stack (circular region of interest around the nucleus). For each worm, a single circular region of interest was also used to measure the background intensity in an adjacent area, and this value was then subtracted from the reporter fluorescence intensity value. For quantification of GFP::CLA-1 and GRASP puncta (**Figure 4**), manual counting was performed using the ImageJ software. For quantification of hypodermal cells (**Figure S5**), fluorescence intensity was measured as described above for individual neurons. For each worm five hypodermal cells were measured, and the fluorescence intensity averaged. The same hypodermal cells were measured in all animals compared.

For fluorescence quantification of CUT rescue lines (**Figure 3**), synchronized day 1 adult worms were grown on NGM plates seeded with OP50 and incubated at 20°C. The COPAS FP-250 system (Union Biometrica; “worm sorter”) was used to measure the fluorescence of 40-150 worms for each strain.

For all behavioral assay, randomization and blinding was done wherever possible. All statistical tests for fluorescence quantifications and behavior assays were conducted using Prism (Graphpad) as described in figure legends.

### SUPPLEMENTAL INFORMATION TITLES AND LEGENDS

**Figure S1.**
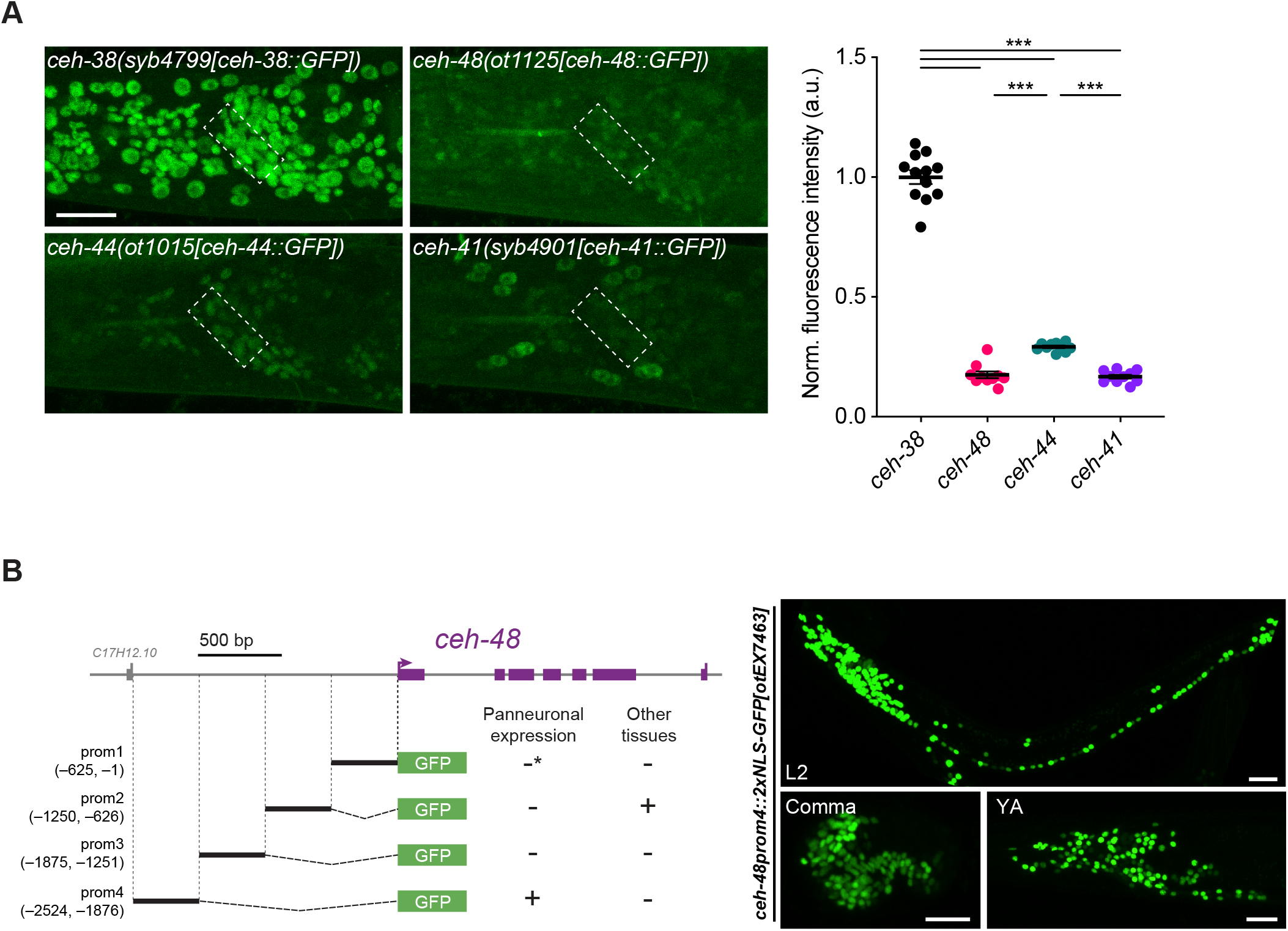
Comparison of the expression level of CUT genes and *ceh-48 cis*- regulatory analysis. (A) Expression of *ceh-38(syb4799[ceh-38::GFP])*, *ceh-48(ot1125[ceh-48::GFP])*, *ceh- 44(ot1015[ceh-44::GFP])* and *ceh-41(syb4901[ceh-41::GFP])* CRISPR/Cas9- engineered *gfp*-tagged alleles. Lateral views of the worm head at the L4 stage are shown. Quantification of reporter allele fluorescence intensity in nerve ring neurons (outlined). The data are presented as individual values with each dot representing the expression level of one worm with the mean ± SEM indicated. One-way ANOVA followed by Tukey’s multiple comparisons test; ***P < 0.001. n ≥ 10 for all genotypes. (B) Schematic representation of *ceh-48* gene loci and *cis-*regulatory analysis. Promoter fragment 4 (prom4) is expressed pan-neuronally and was used to drive pan-neuronal expression in the rescue lines. Promoter fragment 1 (prom1) is expressed in some midbody neurons (asterisk). Promoter fragment 2 (prom2) is expressed in some vulval cells. Expression of a prom4 fusion to GFP (right) is shown at the comma embryonic stage (bottom left, lateral view), L1 larval stage (top, full worm lateral view) and young adult stage (bottom right, lateral view of the head) showing *ceh-48prom4::2xNLS- GFP[otEX7463]* pan-neuronal expression. a.u., arbitrary units. Scale bar 15 μm.

**Figure S2.**
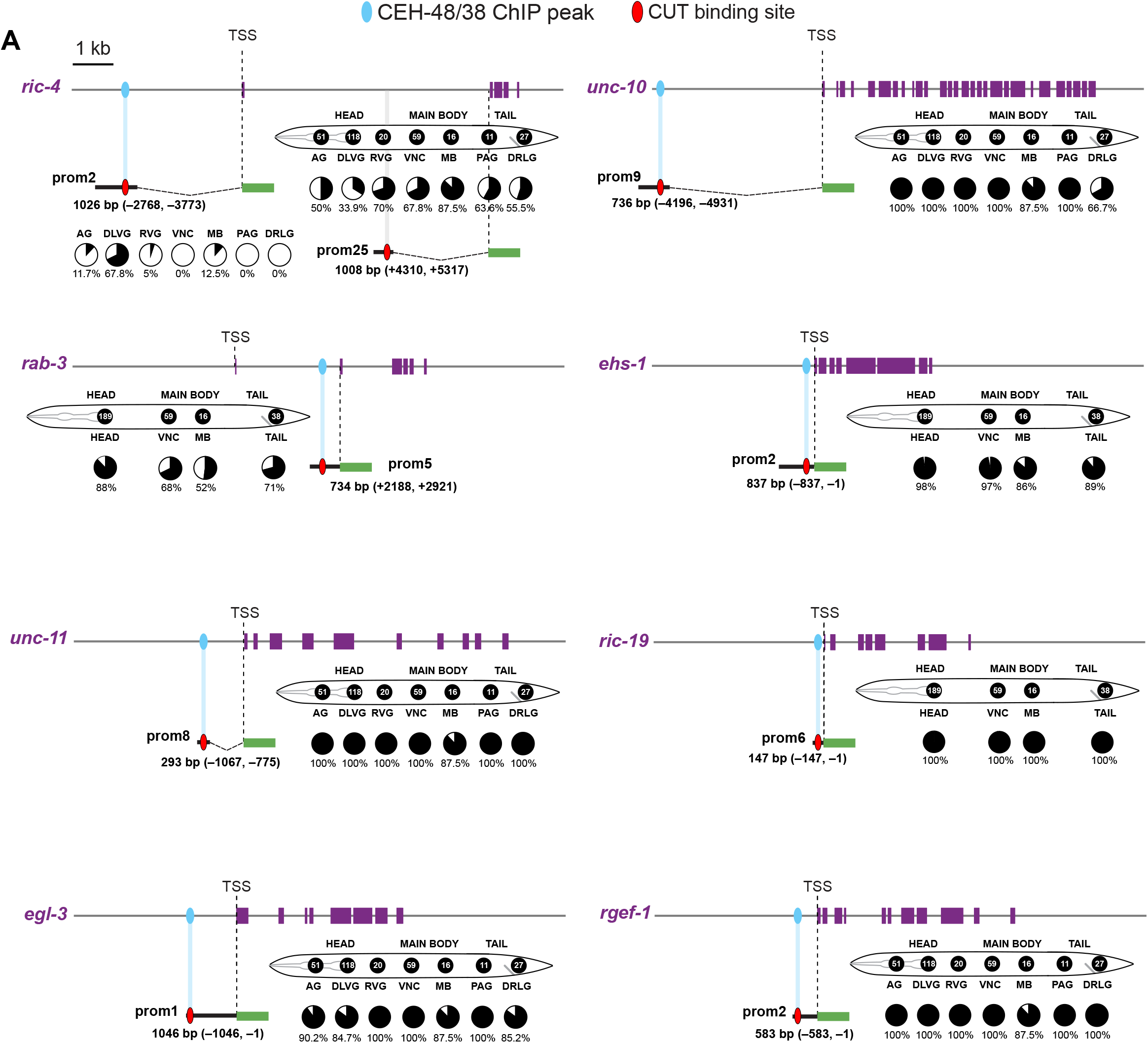
Pan-neuronal *cis*-regulatory elements are bound by CUT factors. (A) CUT ChIP binding correlates with the *cis-*regulatory elements that we previously defined in pan-neuronally expressed genes (Stefanakis et al., 2015). Schematic representation of *ric-4*, *unc-10*, *rab-3*, *ehs-1*, *unc-11*, *ric-19*, *egl-3* and *rgef-1* gene loci. For each gene, a schematic representation of the nervous system of *C. elegans* is shown. Neurons belonging in the different ganglia or regions, are clustered together and represented by a black circle (number of neurons belonging in each ganglion are indicated inside the circle). Below each worm schematic, the fraction of neurons of each ganglion expressing a reporter is indicated with a partially filled circle (pie-chart). Promoter fragment gene fusion reporters expression data from (Stefanakis et al., 2015). For *ric-4*, mutation of the CUT binding site (site 1) overlapping the ChIP peak had no effect on the expression of a *ric-4* CRISPR reporter (Figure 2C). We made use of our previous *ric-4 cis*-regulatory analysis (Stefanakis et al., 2015) to guide our search for CUT binding sites within *cis*-regulatory elements driving broad neuronal expression. We found a cluster of CUT binding sites (site 2) in the previously defined *ric-4* promoter fragment 25 (prom25), located within the first *ric-4* intron. Upon mutation of this cluster, together with site 1, we found a reduction on *ric-4* CRISPR reporter expression (Figure 2C). AG: anterior head ganglion; DLVG: dorsal, lateral, and ventral head ganglia; RVG: retrovesicular ganglion; VNC: ventral nerve cord motor neurons; MB: mid-body neurons; PAG: preanal ganglion; DRLG: dorsorectal and lumbar ganglia. The length in bp and the coordinates of each promoter fragment in relation to the translational start site are shown next to each construct. Promoter fragment number corresponds to the *cis-* regulatory analysis generated in (Stefanakis et al., 2015).

**Figure S3.**
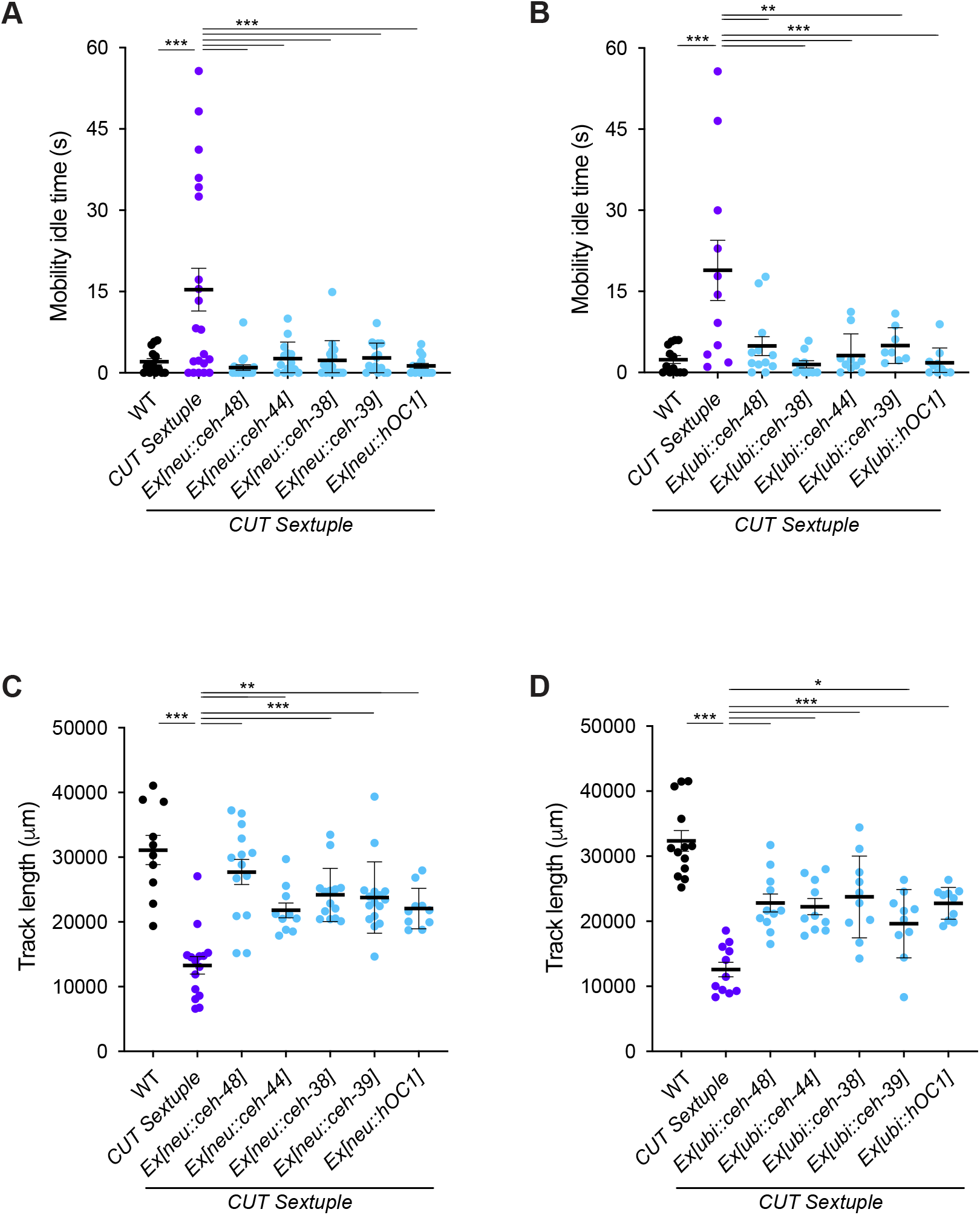
Locomotion phenotypes in CUT sextuple mutant can be rescued by overexpression of individual CUT genes. **(A-D)** Worm mobility idle time (A-B) and track length (C-D) was compared between wild-type, CUT sextuple mutant, and CUT sextuple mutant rescue (panneuronal, *ceh-48* promoter (“neu”) (A and C), or ubiquitous, *eft-3* promoter (“ubi”) (B and D), expression of *ceh-48*, *ceh-44*, *ceh-38*, *ceh-39* or *hOC1*) using a multi-worm tracker system (Roussel et al., 2014). The data are presented as individual values with each dot representing the mobility idle time of one worm with the mean ± SEM indicated. One-way ANOVA followed by Tukey’s multiple comparisons test, comparisons with CUT sextuple mutant indicated; **P < 0.01, ***P < 0.001. n ≥ 10 for all genotypes. WT, wild-type

**Figure S4.**
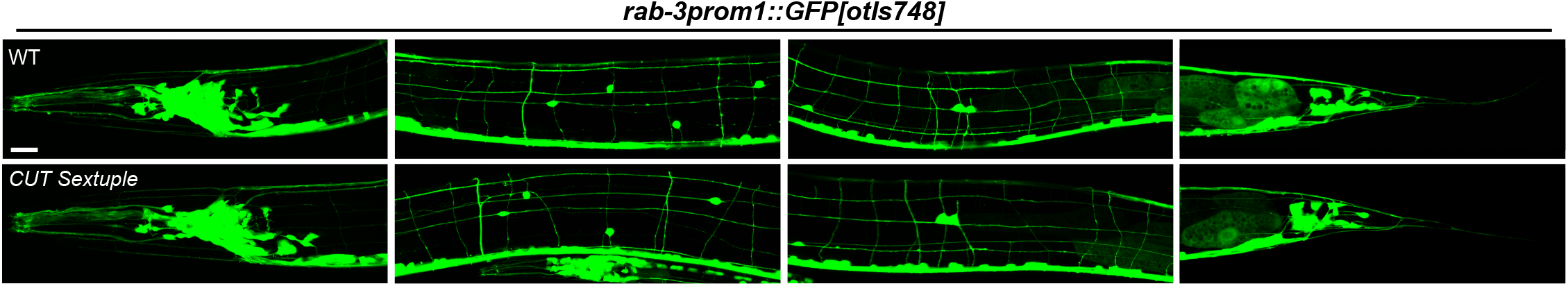
Overall neuronal anatomy is not affected in CUT gene mutant. Overall nervous system anatomy (visualized through a *rab-3* cytoplasmic reporter, *rab- 3prom1::GFP[otIs748]*) is not affected in CUT sextuple mutant (bottom) when compared to wild-type (top) animals. All images correspond to worms at the L4 larval stage. Signal is shown saturated in both genotypes to visualize the axonal tracts, therefore the defect on *rab-3* expression in the CUT sextuple mutant is not visible here. WT, wild-type; Scale bar 15 μm.

**Figure S5.**
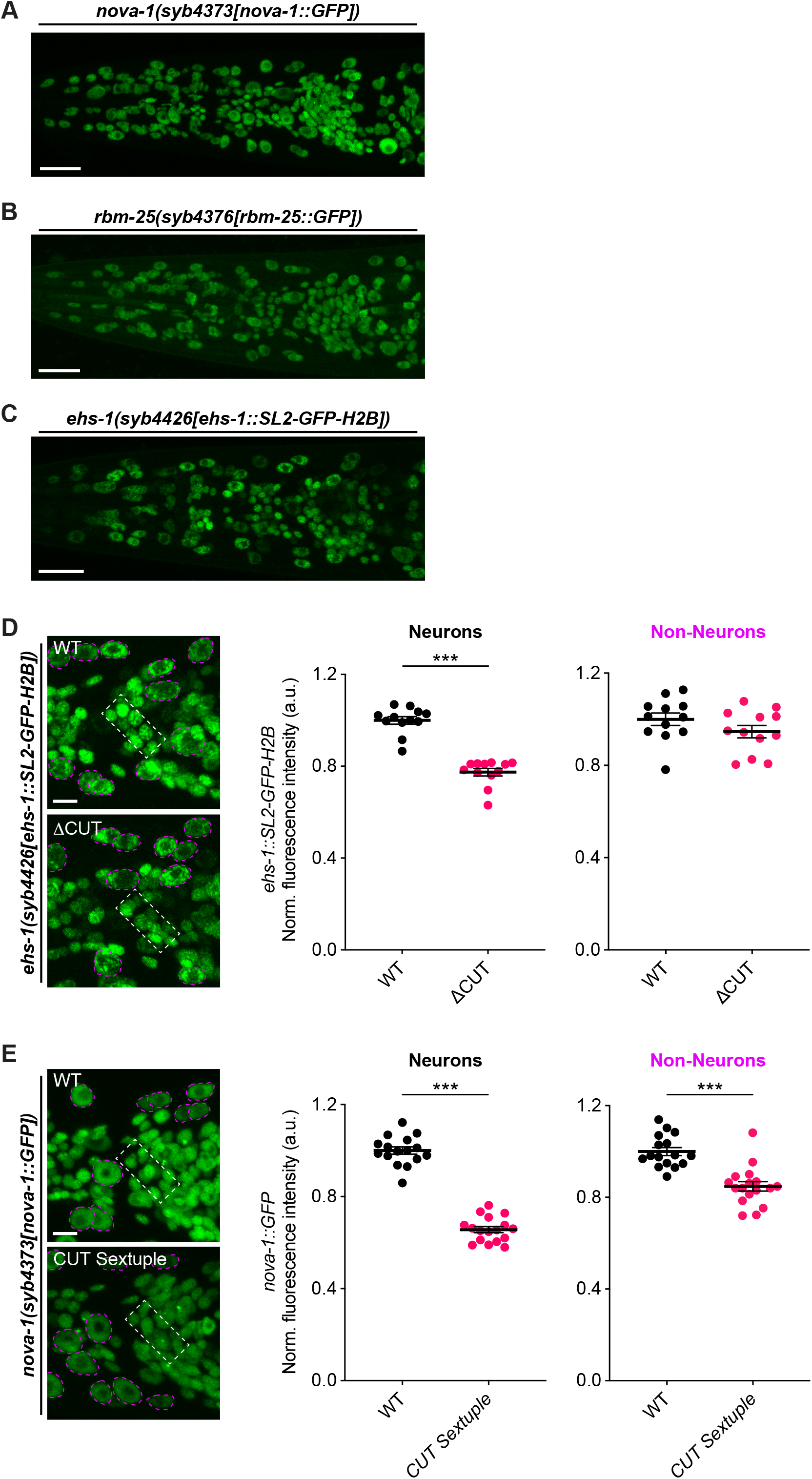
CUT genes regulate some ubiquitously expressed genes with neuronal functions. (A-C) Expression of nova-1(syb4373[nova-1::GFP]) (A), rbm-25(syb4376[rbm-25::GFP]) (B) and ehs-1(syb4426[ehs-1::SL2-GFP-H2B]) (C) CRISPR/Cas9-engineered gfp-tagged alleles in wild-type animals. These genes are downregulated in the CUT sextuple mutant transcriptional analysis (ehs-1 q-value is just above the significant threshold, q-value = 0.070, p-value = 0.015). (D) CUT binding site mutation in the ehs-1 endogenous promoter (bottom) causes a reduction on ehs-1 expression in neurons compared to wild-type (top), but not in non-neuronal cells. Lateral views of the worm head at the L4 stage are shown. Quantification of ehs-1(syb4426[ehs-1::SL2-GFP-H2B]) fluorescence intensity in nerve ring neurons (white outlined box) and head epidermal cells (outlined in magenta). The data are presented as individual values with each dot representing the expression level of one worm with the mean ± SEM indicated. Unpaired t-test, ***P < 0.001. n = 12 for all genotypes. (E) Expression of nova-1(syb4373[nova-1::GFP]) in wild-type (top) and CUT sextuple mutant (bottom). Lateral views of the worm head at the L4 stage are shown. Quantification of nova-1(syb4373[nova-1::GFP]) fluorescence intensity in nerve ring neurons (white outlined box) and head epidermal cells (outlined in magenta). The data are presented as individual values with each dot representing the expression level of one worm with the mean ± SEM indicated. Unpaired t-test, ***P < 0.001. n ≥ 16 for all genotypes WT, wild-type; a.u., arbitrary units. Scale bars 15 μm (A-C), 5 μm (D-E).

**Table S1 Genes associated to CEH-48 and CEH-38 ChIP datasets**

CEH-48 (Experiment: ENCSR844VCY, https://www.encodeproject.org/) and CEH-38 (Accession # modEncode_4800, http://www.modencode.org/) ChIP-seq datasets were used to identify genes associated to the peak regions. Those genes which promoter region (defined as from 5kb upstream of the transcription start site to 1kb downstream) overlapped with the ChIP-seq peaks were identified. A list of genes associated to CEH- 48 peaks (4709 genes), genes associated to CEH-38 peaks (7795 genes), and genes associated to CEH-48 or CEH-38 peaks (9578 genes) are provided in separate tabs. The genes are sorted by WormBase Gene ID. Brief gene descriptions obtained from WormMine.

**Table S2 Differentially expressed genes in CUT sextuple mutant**

Comparison of neuronal-nuclei immunoprecipitated samples from wild-type and CUT sextuple mutant is conducted using DESeq2 (Love et al., 2014) to determine differentially expressed genes (DEG, 3834 genes have a padj < 0.05). The genes are sorted by padj from smallest to largest. baseMean is the average normalized read counts of all samples. log2FoldChange is calculated using the formula log2 (average read counts of mutant samples)/(average read counts of wild-type samples). A list of downregulated genes (genes with a log2FoldChange < 0; 2118 genes), upregulated genes (genes with a log2FoldChange > 0; 1716 genes), and differentially expressed genes that are found associated to CEH-48 or CEH-38 ChIP-seq peaks (1124 genes) are provided in separate tabs, with the genes sorted by WormBase Gene ID. Brief gene descriptions obtained from WormMine.

**Table S3 Neuronally-enriched genes**

Comparison of wild-type neuronal-nuclei immunoprecipitated (IP) samples to wild-type input (total nuclei) samples is conducted using DESeq2 to determine enrichment (12639 genes have a padj < 0.05). The genes are sorted by padj from smallest to largest. baseMean is the average normalized read counts of all IP and input samples. log2FoldChange is calculated using the formula log2 (average read counts of mutant samples)/(average read counts of wild-type samples). A list of neuronally-enriched genes (genes with a log2FoldChange > 0; 6372 genes), and neuronally-depleted genes (genes with a log2FoldChange < 0; 6267 genes) are provided in separate tabs, with the genes sorted by WormBase Gene ID. Brief gene descriptions obtained from WormMine.

**Table S4 Full list of enrichment analysis GO and phenotype terms**

Full list of enriched terms for GO enrichment analysis and phenotype enrichment analysis using gene sets of significantly downregulated or upregulated transcripts. The terms are sorted by q-value from smallest to largest. Analysis performed using the Gene Set Enrichment Analysis Tool from Wormbase.

**Table S5 Known transcription factor motifs matching CEH-48 and CEH-38 motifs**

Tomtom Motif Comparison Tool (Gupta et al., 2007) output list of known motifs matching the CEH-48 motif defined in (Weirauch et al., 2014), the CEH-48 motif extracted from the ChIP-seq dataset (Experiment: ENCSR844VCY, https://www.encodeproject.org/) using MEME-ChIP (Machanick and Bailey, 2011), and the CEH-38 motif extracted from the ChIP-seq dataset (Accession # modEncode_4800, http://www.modencode.org/) using MEME-ChIP. These three motifs were compared with an eukaryote DNA motif database (vertebrates, in vivo and in silico) provided in MEME Suite (Gupta et al., 2007) (JASPAR2018_CORE_vertebrates_non-redundant, uniprobe_mouse, and jolma2013; total 1808 known motifs). The motifs are sorted by q-value from smallest to largest.

## REFERENCES

Alkema, M.J., Hunter-Ensor, M., Ringstad, N., and Horvitz, H.R. (2005). Tyramine Functions independently of octopamine in the Caenorhabditis elegans nervous system. Neuron 46, 247–260.

Altun-Gultekin, Z., Andachi, Y., Tsalik, E.L., Pilgrim, D., Kohara, Y., and Hobert, O. (2001). A regulatory cascade of three homeobox genes, ceh-10, ttx-3 and ceh-23, controls cell fate specification of a defined interneuron class in C. elegans. Development 128, 1951–1969.

Bertrand, N., Castro, D.S., and Guillemot, F. (2002). Proneural genes and the specification of neural cell types. Nat Rev Neurosci 3, 517–530.

Brenner, S. (1974). The genetics of Caenorhabditis elegans. Genetics 77, 71–94.

Burglin, T.R., and Affolter, M. (2016). Homeodomain proteins: an update. Chromosoma 125, 497–521.

Burglin, T.R., and Cassata, G. (2002). Loss and gain of domains during evolution of cut superclass homeobox genes. Int J Dev Biol 46, 115–123.

Cinar, H., Keles, S., and Jin, Y. (2005). Expression profiling of GABAergic motor neurons in Caenorhabditis elegans. Curr Biol 15, 340–346.

Croll, N.A. (1975). Behavioural analysis of nematode movement. Adv Parasitol 13, 71–122.

Davis, C.A., Hitz, B.C., Sloan, C.A., Chan, E.T., Davidson, J.M., Gabdank, I., Hilton, J.A., Jain, K., Baymuradov, U.K., Narayanan, A.K., et al. (2018). The Encyclopedia of DNA elements (ENCODE): data portal update. Nucleic Acids Res 46, D794–D801.

Dobin, A., Davis, C.A., Schlesinger, F., Drenkow, J., Zaleski, C., Jha, S., Batut, P., Chaisson, M., and Gingeras, T.R. (2013). STAR: ultrafast universal RNA-seq aligner. Bioinformatics 29, 15–21.

Dokshin, G.A., Ghanta, K.S., Piscopo, K.M., and Mello, C.C. (2018). Robust Genome Editing with Short Single-Stranded and Long, Partially Single-Stranded DNA Donors in Caenorhabditis elegans. Genetics 210, 781–787.

Dykes, I.M., Tempest, L., Lee, S.I., and Turner, E.E. (2011). Brn3a and Islet1 act epistatically to regulate the gene expression program of sensory differentiation. J Neurosci 31, 9789–9799.

Feinberg, E.H., Vanhoven, M.K., Bendesky, A., Wang, G., Fetter, R.D., Shen, K., and Bargmann, C.I. (2008). GFP Reconstitution Across Synaptic Partners (GRASP) defines cell contacts and synapses in living nervous systems. Neuron 57, 353–363.

Gupta, S., Stamatoyannopoulos, J.A., Bailey, T.L., and Noble, W.S. (2007). Quantifying similarity between motifs. Genome Biol 8, R24.

Hobert, O. (2002). PCR fusion-based approach to create reporter gene constructs for expression analysis in transgenic C. elegans. BioTechniques 32, 728–730.

Hobert, O. (2016). Terminal Selectors of Neuronal Identity. Curr Top Dev Biol 116, 455–475.

Hobert, O. (2021). Homeobox genes and the specification of neuronal identity. Nat Rev Neurosci 22, 627–636.

Hobert, O., Carrera, I., and Stefanakis, N. (2010). The molecular and gene regulatory signature of a neuron. Trends in neurosciences 33, 435–445.

Hobert, O., and Kratsios, P. (2019). Neuronal identity control by terminal selectors in worms, flies, and chordates. Curr Opin Neurobiol 56, 97–105.

Ioannou, M.S., and Marat, A.L. (2012). The role of EHD proteins at the neuronal synapse. Sci Signal 5, jc1.

Jensen, K.B., Dredge, B.K., Stefani, G., Zhong, R., Buckanovich, R.J., Okano, H.J., Yang, Y.Y., and Darnell, R.B. (2000). Nova-1 regulates neuron-specific alternative splicing and is essential for neuronal viability. Neuron 25, 359–371.

Kagoshima, H., Cassata, G., Tong, Y.G., Pujol, N., Niklaus, G., and Burglin, T.R. (2013). The LIM homeobox gene ceh-14 is required for phasmid function and neurite outgrowth. Dev Biol 380, 314–323.

Kohn, R.E., Duerr, J.S., McManus, J.R., Duke, A., Rakow, T.L., Maruyama, H., Moulder, G., Maruyama, I.N., Barstead, R.J., and Rand, J.B. (2000). Expression of multiple UNC- 13 proteins in the Caenorhabditis elegans nervous system. Mol Biol Cell 11, 3441–3452.

Kratsios, P., Stolfi, A., Levine, M., and Hobert, O. (2011). Coordinated regulation of cholinergic motor neuron traits through a conserved terminal selector gene. Nat Neurosci 15, 205–214.

Kropp, P.A., and Gannon, M. (2016). Onecut transcription factors in development and disease. Trends Dev Biol 9, 43–57.

Leyva-Diaz, E., Masoudi, N., Serrano-Saiz, E., Glenwinkel, L., and Hobert, O. (2020). Brn3/POU-IV-type POU homeobox genes-Paradigmatic regulators of neuronal identity across phylogeny. Wiley interdisciplinary reviews Developmental biology, e374.

Leyva-Diaz, E., Stefanakis, N., Carrera, I., Glenwinkel, L., Wang, G., Driscoll, M., and Hobert, O. (2017). Silencing of Repetitive DNA Is Controlled by a Member of an Unusual Caenorhabditis elegans Gene Family. Genetics 207, 529–545.

Liao, Y., Smyth, G.K., and Shi, W. (2014). featureCounts: an efficient general purpose program for assigning sequence reads to genomic features. Bioinformatics 30, 923–930.

Lloret-Fernandez, C., Maicas, M., Mora-Martinez, C., Artacho, A., Jimeno-Martin, A., Chirivella, L., Weinberg, P., and Flames, N. (2018). A transcription factor collective defines the HSN serotonergic neuron regulatory landscape. eLife 7.

Love, M.I., Huber, W., and Anders, S. (2014). Moderated estimation of fold change and dispersion for RNA-seq data with DESeq2. Genome Biol 15, 550.

Lowe, E.K., and Stolfi, A. (2018). Developmental system drift in motor ganglion patterning between distantly related tunicates. Evodevo 9, 18.

Machanick, P., and Bailey, T.L. (2011). MEME-ChIP: motif analysis of large DNA datasets. Bioinformatics 27, 1696–1697.

Mahoney, T.R., Luo, S., and Nonet, M.L. (2006). Analysis of synaptic transmission in Caenorhabditis elegans using an aldicarb-sensitivity assay. Nat Protoc 1, 1772–1777.

Nguyen, D.N., Rohrbaugh, M., and Lai, Z. (2000). The Drosophila homolog of Onecut homeodomain proteins is a neural-specific transcriptional activator with a potential role in regulating neural differentiation. Mech Dev 97, 57–72.

Nguyen, M., Alfonso, A., Johnson, C.D., and Rand, J.B. (1995). Caenorhabditis elegans mutants resistant to inhibitors of acetylcholinesterase. Genetics 140, 527–535.

Nonet, M.L., Staunton, J.E., Kilgard, M.P., Fergestad, T., Hartwieg, E., Horvitz, H.R., Jorgensen, E.M., and Meyer, B.J. (1997). Caenorhabditis elegans rab-3 mutant synapses exhibit impaired function and are partially depleted of vesicles. J Neurosci 17, 8061–8073.

Pereira, L., Kratsios, P., Serrano-Saiz, E., Sheftel, H., Mayo, A.E., Hall, D.H., White, J.G., LeBoeuf, B., Garcia, L.R., Alon, U., et al. (2015). A cellular and regulatory map of the cholinergic nervous system of C. elegans. eLife 4.

Pierce, M.L., Weston, M.D., Fritzsch, B., Gabel, H.W., Ruvkun, G., and Soukup, G.A. (2008). MicroRNA-183 family conservation and ciliated neurosensory organ expression. Evol Dev 10, 106–113.

Pierce-Shimomura, J.T., Chen, B.L., Mun, J.J., Ho, R., Sarkis, R., and McIntire, S.L. (2008). Genetic analysis of crawling and swimming locomotory patterns in C. elegans. Proc Natl Acad Sci U S A 105, 20982–20987.

Poustka, A.J., Kuhn, A., Radosavljevic, V., Wellenreuther, R., Lehrach, H., and Panopoulou, G. (2004). On the origin of the chordate central nervous system: expression of onecut in the sea urchin embryo. Evol Dev 6, 227–236.

Qiu, Z., and Ghosh, A. (2008). A brief history of neuronal gene expression: regulatory mechanisms and cellular consequences. Neuron 60, 449–455.

Reilly, M.B., Cros, C., Varol, E., Yemini, E., and Hobert, O. (2020). Unique homeobox codes delineate all the neuron classes of C. elegans. Nature 584, 595–601.

Restif, C., Ibanez-Ventoso, C., Vora, M.M., Guo, S., Metaxas, D., and Driscoll, M. (2014). CeleST: computer vision software for quantitative analysis of C. elegans swim behavior reveals novel features of locomotion. PLoS Comput Biol 10, e1003702.

Roussel, N., Sprenger, J., Tappan, S.J., and Glaser, J.R. (2014). Robust tracking and quantification of C. elegans body shape and locomotion through coiling, entanglement, and omega bends. Worm 3, e982437.

Schneider, C.A., Rasband, W.S., and Eliceiri, K.W. (2012). NIH Image to ImageJ: 25 years of image analysis. Nat Methods 9, 671–675.

Serrano-Saiz, E., Gulez, B., Pereira, L., Gendrel, M., Kerk, S.Y., Vidal, B., Feng, W., Wang, C., Kratsios, P., Rand, J.B., et al. (2020). Modular Organization of Cis-regulatory Control Information of Neurotransmitter Pathway Genes in Caenorhabditis elegans. Genetics 215, 665–681.

Serrano-Saiz, E., Poole, Richard J., Felton, T., Zhang, F., De La Cruz, Estanisla D., and Hobert, O. (2013). Modular Control of Glutamatergic Neuronal Identity in C. elegans by Distinct Homeodomain Proteins. Cell 155, 659–673.

Stefanakis, N., Carrera, I., and Hobert, O. (2015). Regulatory Logic of Pan-Neuronal Gene Expression in C. elegans. Neuron 87, 733–750.

Steiner, F.A., Talbert, P.B., Kasinathan, S., Deal, R.B., and Henikoff, S. (2012). Cell-type-specific nuclei purification from whole animals for genome-wide expression and chromatin profiling. Genome Res 22, 766–777.

Sun, H., and Hobert, O. (2021). Temporal transitions in the post-mitotic nervous system of Caenorhabditis elegans. Nature 600, 93–99.

Taylor, S.R., Santpere, G., Weinreb, A., Barrett, A., Reilly, M.B., Xu, C., Varol, E., Oikonomou, P., Glenwinkel, L., McWhirter, R., et al. (2021). Molecular topography of an entire nervous system. Cell 184, 4329–4347 e4323.

Vassalli, Q.A., Colantuono, C., Nittoli, V., Ferraioli, A., Fasano, G., Berruto, F., Chiusano, M.L., Kelsh, R.N., Sordino, P., and Locascio, A. (2021). Onecut Regulates Core Components of the Molecular Machinery for Neurotransmission in Photoreceptor Differentiation. Front Cell Dev Biol 9, 602450.

Weirauch, M.T., Yang, A., Albu, M., Cote, A.G., Montenegro-Montero, A., Drewe, P., Najafabadi, H.S., Lambert, S.A., Mann, I., Cook, K., et al. (2014). Determination and inference of eukaryotic transcription factor sequence specificity. Cell 158, 1431–1443.

Weiss, L.A., and Nieto, M. (2019). The crux of Cux genes in neuronal function and plasticity. Brain Res 1705, 32–42.

Yemini, E., Jucikas, T., Grundy, L.J., Brown, A.E., and Schafer, W.R. (2013). A database of Caenorhabditis elegans behavioral phenotypes. Nat Methods 10, 877–879.

Yu, B., Wang, X., Wei, S., Fu, T., Dzakah, E.E., Waqas, A., Walthall, W.W., and Shan, G. (2017). Convergent Transcriptional Programs Regulate cAMP Levels in C. elegans GABAergic Motor Neurons. Dev Cell 43, 212–226 e217.

